# Gene expression drives the evolution of dominance

**DOI:** 10.1101/182865

**Authors:** Christian D. Huber, Arun Durvasula, Angela M. Hancock, Kirk E. Lohmueller

## Abstract

Dominance is a fundamental concept in molecular genetics and has implications for understanding patterns of genetic variation, evolution, and complex traits. However, despite its importance, the degree of dominance has yet to be quantified in natural populations. Here, we leverage multiple mating systems in natural populations of *Arabidopsis* to co-estimate the distribution of fitness effects and dominance coefficients of new amino acid changing mutations. We find that more deleterious mutations are more likely to be recessive than less deleterious mutations. Further, this pattern holds across gene categories, but varies with the connectivity and expression patterns of genes. Our work argues that dominance arose as the inevitable consequence of the functional importance of genes and their optimal expression levels.

**One sentence summary:** We use population genomic data to characterize the degree of dominance for new mutations and develop a new theory for its evolution.

---

The relationship between the fitness effects of heterozygous and homozygous genotypes at a locus, termed dominance, is the major factor that determines the fate of new alleles in a population and has far reaching implications for genetic diseases and evolutionary genetics (1-4). Several models have been theorized for the mechanism of dominance, starting with R.A. Fisher's model, which suggests that dominance arises via modifier mutations at other loci and that these loci are subject to selection (5). In response, S. Wright argued that selection would not be strong enough to maintain these modifier mutations. He proposed a different model (termed the “metabolic theory”), later extended by Kacser and Burns, predicting most mutations in enzymes will be recessive because the reduced activity of mutant alleles can be masked by the wild type allele in heterozygotes (6, 7). An alternative model, posited by Haldane and further developed by Hurst and Randerson, suggested that recessivity is a consequence of selection for higher amounts of enzyme product because enzymes expressed at higher levels are able to tolerate loss of function (LoF) mutations (8, 9).

The Wright and Haldane models predict that there is a negative relationship between the dominance coefficient (h) and the selection coefficient (s), such that more deleterious mutations will tend to be recessive, while Fisher’s model makes no such prediction. Drosophila mutation accumulation lines showed evidence of this relationship, providing the first empirical evidence that Fisher's theory may not hold (10-12). While the predictions of the Wright and Haldane models may be applicable to enzymes, they fail to explain the mechanism of dominance in noncatalytic gene products (13). Further, the extent to which these estimates apply to the majority of mutations occurring in natural populations remains to be tested. While population genetic approaches to estimate the degree of dominance from segregating genetic variation exist (14, 15), they have not been widely applied to empirical data nor have they been used to test models regarding the evolution of dominance.

A major challenge to studying dominance in natural populations is that *h* is inherently confounded with the distribution of fitness effects (DFE) such that different values of *h* and DFEs can yield similar patterns in the genetic variation data in a single outcrossing population (Fig. 1A). However, these same models can be distinguished from each other by studying organisms that undergo self-fertilization as selection will have the chance to immediately act on recessive homozygotes (Fig. 1B). Here, we leverage this fact by developing a composite likelihood approach, which uses the site frequency spectrum (SFS) of the outcrossing *A. lyrata* and the selfing *A. thaliana* (Fig. 1C) to co-estimate the DFE and distribution of *h* for new nonsynonymous mutations on recently published datasets from both species (16-18).

**Fig 1.**
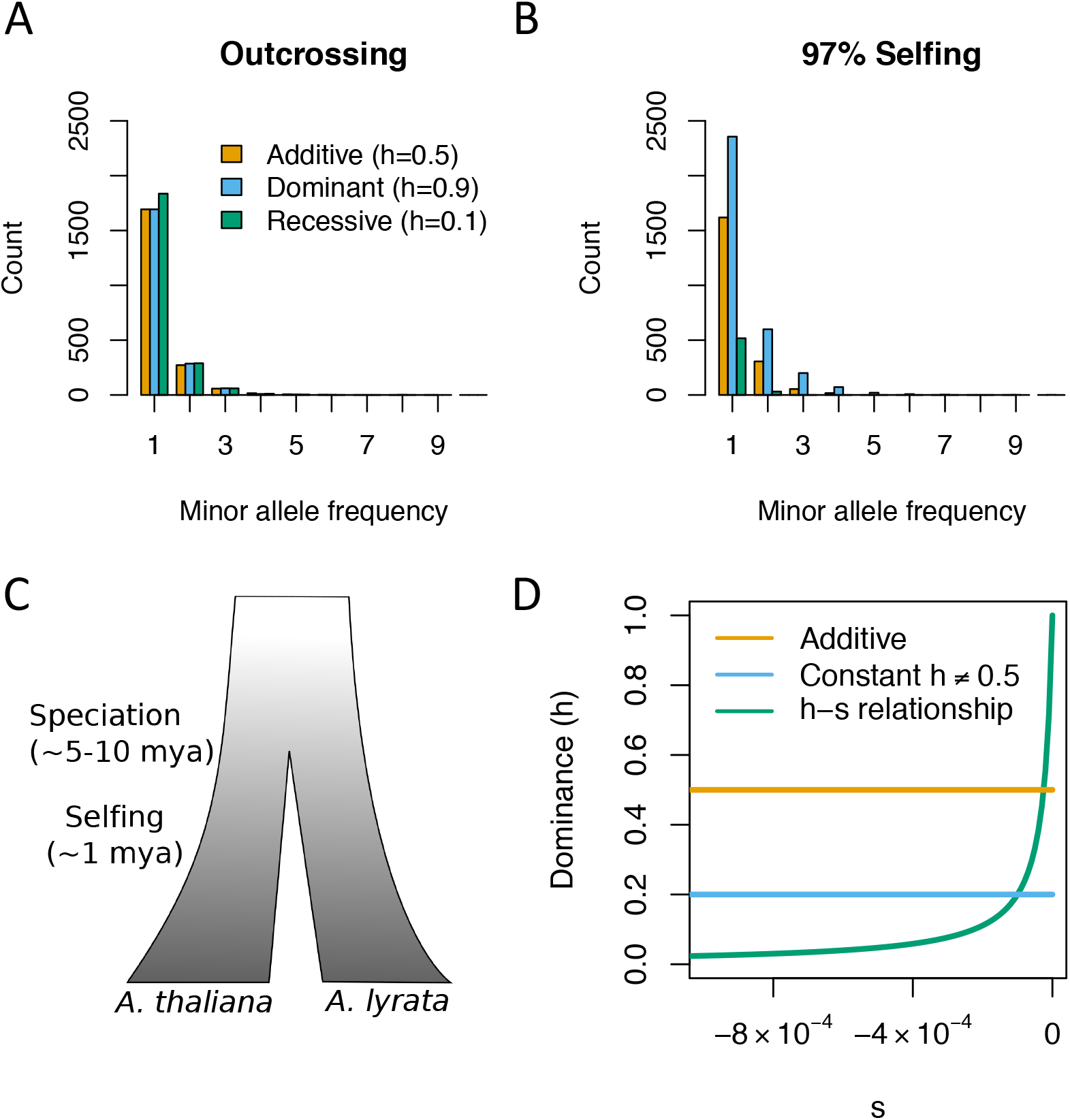
The effect of dominance and mating system on the site frequency spectrum (SFS). **(A)** The SFS from an outcrossing species simulated under different DFEs and *h* values. Note that different combinations of DFEs and values of *h* yield similar SFS. **(B)** The SFS for the same DFEs and values of *h* as in **(A)** for a highly selfing species. Differences in *h* result in large differences in the SFS in selfing species, allowing us to reliably co-estimate the DFE and *h*. **(C)** A schematic of the species history between *A. thaliana* and *A. lyrata*. **(D)** Examples of the relationship between *h* and s under the three different models of dominance tested here.

We compare the fit of 3 distinct models of dominance effects to the SFS from both populations of *Arabidopsis* (Fig. 1D). We find that a model where mutations are slightly recessive (inferred h=0.46) results in a significantly better fit than assuming a model where all mutations are additive (Fig. 2A). The third model allows *h* to depend on s (Fig. 1D), and we infer that this model fits the SFS significantly better than a model with a constant *h* (P<1 × 10^-15^; Fig. 2A) (18). Importantly, mutations that are more deleterious also tend to be more recessive (Fig. 2B). For example, mutations with s<-0.001 have an h<0.025, suggesting that even moderately deleterious mutations are quite recessive. However, because very strongly deleterious mutations (s<-0.01) are unlikely to be segregating in the data, we have limited resolution to infer the dominance effects for such mutations.

**Fig 2.**
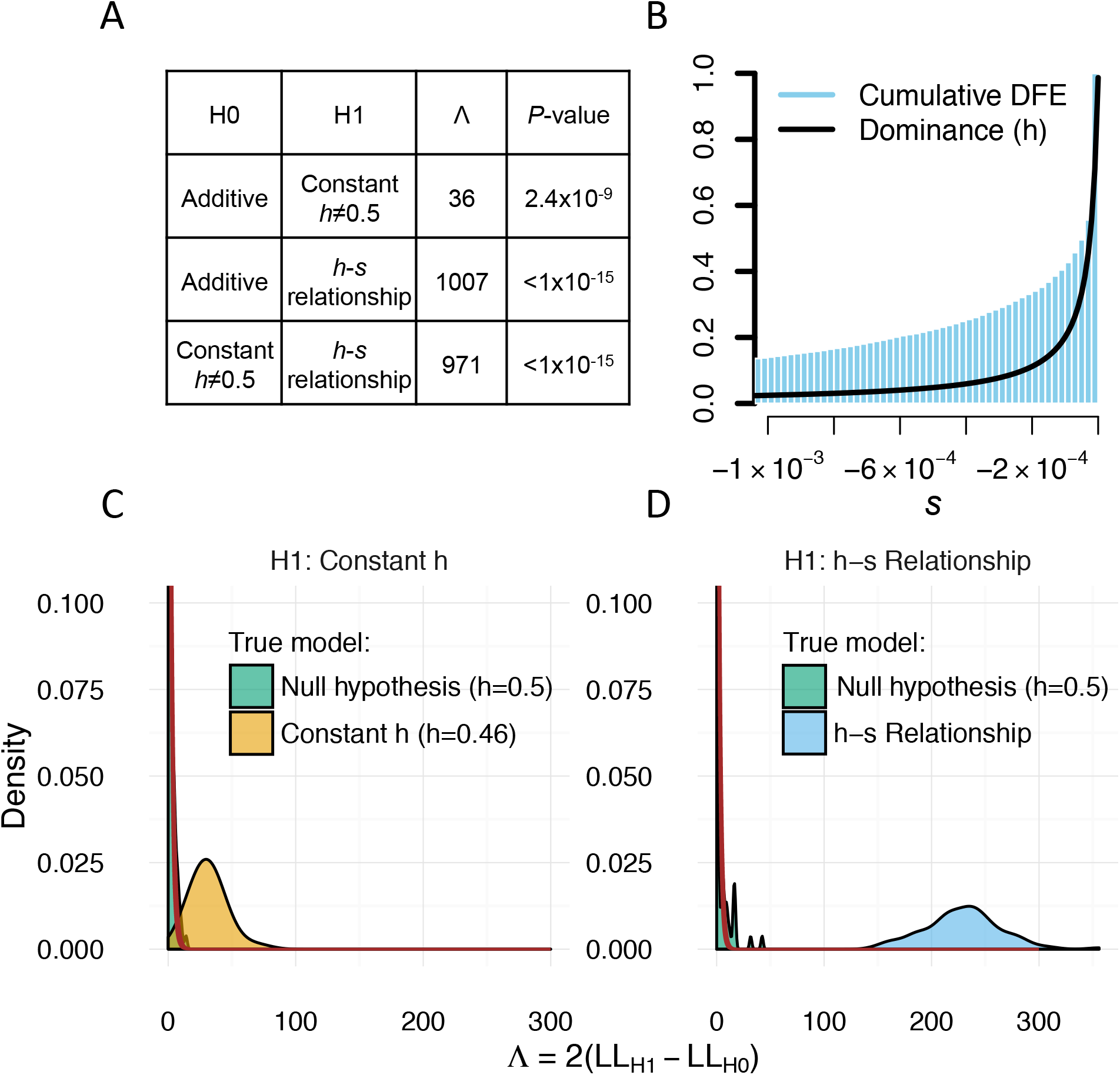
Genome-wide estimates of dominance. **(A)** Likelihood ratio test statistics *(Λ)* and *P*-values when comparing different models of dominance. The *h-s* relationship fits the data significantly better than the additive model and significantly better than a model with a single dominance coefficient. **(B)** Inferred relationship between *h* and *s* based on whole genome data. More nearly neutral mutations tend to be more dominant than strongly deleterious mutations. **(C, D)** Simulations demonstrating the performance of our inference procedure. **(C)** Likelihood ratio tests comparing a constant *h* model to an additive model. When data are simulated under an additive model (green), *Λ* nearly follows a chi-square (1 *df)* distribution (red line). However, when the data are simulated under a model with h=0.46 (tan), the distribution of *Λ* is substantially larger, indicating excellent statistical power. **(D)** Likelihood ratio tests comparing the *h-s* relationship model to an additive model. When data are simulated under an additive model (green), *Λ* nearly follows a chi-square (2 *df*) distribution (red line). However, when the data are simulated under the *h-s* relationship model (tan), the distribution of *Λ* is substantially larger, indicating excellent statistical power.

To determine whether our statistical framework is sensitive to certain confounders and can reliably distinguish between competing models, we carried out extensive forward simulations based on the demographic models inferred from our data (table S1, fig. S1)(18). The distribution of the likelihood ratio test (LRT) statistic in simulations where all mutations were additive resembled the predicted asymptotic chi-square distribution when comparing the constant *h≠0.5* model to the additive model (df=1, Fig. 2C) as well as when comparing the *h-s* relationship model to the additive model (df=2, Fig. 2D). Importantly, none of the LRT statistics were as large as those seen empirically (Fig. 2A), suggesting a conservative simulation-based P-value <0.01. When simulating data under the constant *h* model (h=0.46, Fig. 2C) as well as the *h-s* relationship model, we find that the distribution of the LRT statistic is much greater than that of the null data (Fig. 2D). These simulations suggest we have excellent power to distinguish between models given the demographic history, sample size, and amounts of genetic variation present in these species. Lastly, our simulations show that differing DFEs between *A. thaliana* and *A. lyrata* would not provide false evidence of the *h-s* relationship (18) (fig. S2, tables S2 and S3). In sum, it is unlikely that our conclusion of extensive recessivity of mutations and the relationship between dominance effects and selective effects is driven by artifacts of our inference procedure.

We next sought to test which theoretical model for the evolution of dominance can explain our data. Fisher’s theory for the evolution of dominance predicts that *h* should show no relationship to the degree of deleteriousness of a mutation (5). Our finding of the *h-s* relationship is not consistent with this theory. The metabolic theory (7) predicts that mutations in catalytic genes ought to be more recessive than those in genes unlikely to be involved in enzyme kinetics. We classified genes based on Gene Ontology (GO) category and inferred the DFE and *h* on specific gene sets (18) (tables S3 and S4). Overall, we find that catalytic genes display similar patterns of polymorphism (fig. S4) and an *h-s* relationship, as seen genome-wide (Fig. 3A). Genes encoding structural proteins (herein “structural genes”), which are unlikely to be involved in enzyme kinetics, however, show a higher proportion of rare variants in the SFS (fig. S4) and appear to be less recessive than catalytic genes (Fig. 3A). In other words, for a given selection coefficient, mutations in catalytic genes tend to be more recessive than those in structural genes. On the surface, this finding appears to support the prediction of the metabolic theory of dominance. However, we infer that the *h-s* relationship model fits the structural genes better than the constant *h* model or the additive model (Fig. 3C, table S3). Thus, even structural genes show evidence of recessive mutations, which is not predicted under the metabolic theory model. We note that this finding has previous experimental support in yeast (13, 19).

**Fig 3.**
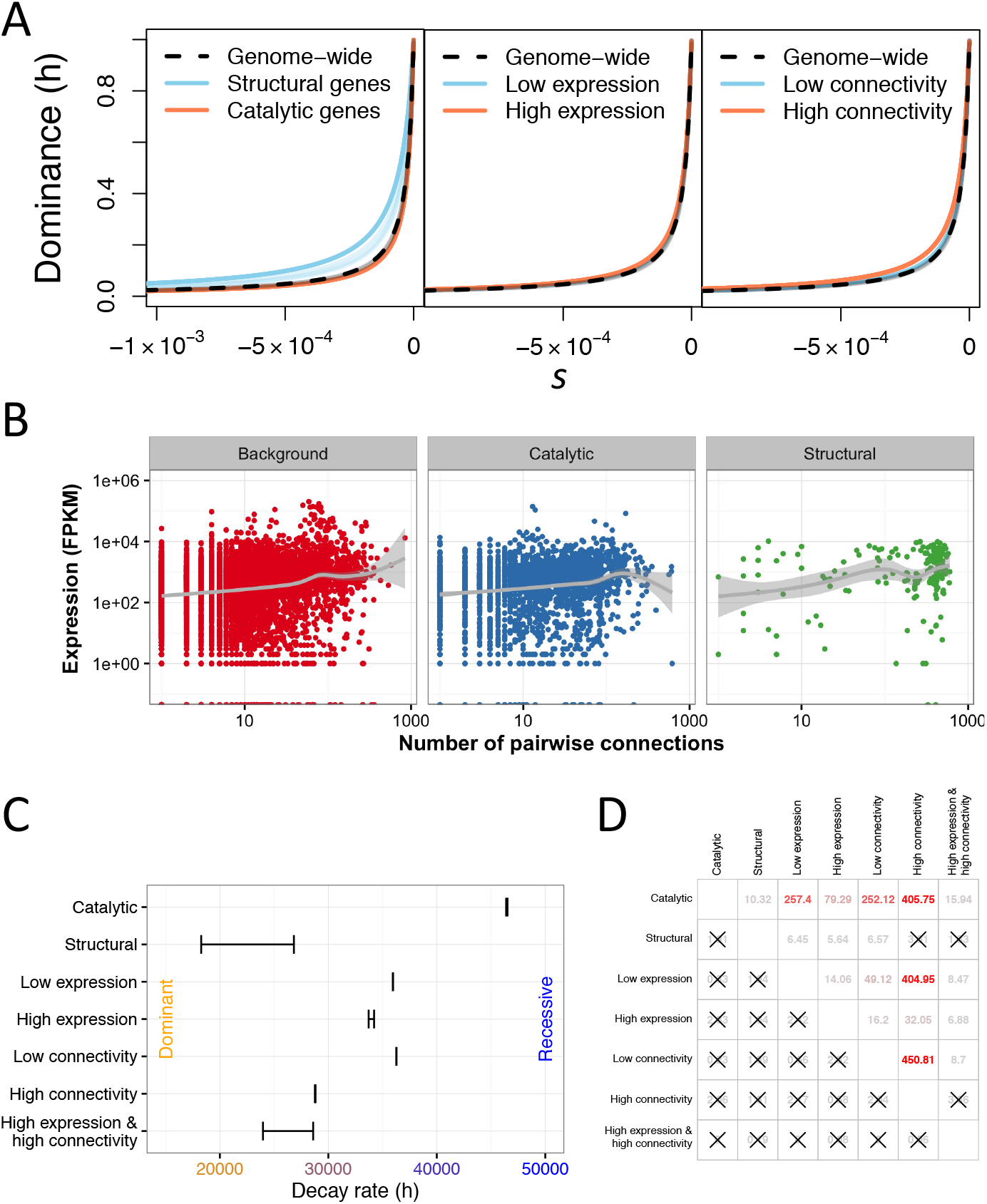
Distribution of dominance per gene category. **(A)** *h-s* relationship inferred for different gene categories. Bootstrap replicates are shown in lighter colors. **(B)** Expression profiles are correlated with gene connectivity. Note that structural genes have higher connectivity and expression than do other types of genes. Background refers to genes not in catalytic or structural GO categories. **(C)** Differences in the decay rate of *h* across gene categories. 95% confidence intervals (CI) are shown. Larger decay rates indicate that for a given value of s, mutations tend to be more recessive. **(D)** Z-scores for tests of differences in decay rate (upper triangle) and intercept (lower triangle) between different categories of genes. Color indicates degree of significance (red is more significant). Comparisons not significantly different after Bonferroni correction are denoted by “X”s.

To investigate other mechanisms that could lead to recessive mutations in structural genes, we classified genes based on their expression level and degree of connectivity in networks (18). Overall, we found that structural genes tended to be more highly expressed and have more network connections than other types of genes (Fig. 3B). We next tested whether the parameters of the *h-s* relationship differed across these different functional categories (Fig. 3C and 3D, figs. S7 and S8, tables S3 and S4). While the *h* intercept did not differ across any of the categories (Fig. 3D, fig. S7), we found that the *h-s* decay rate, or slope, of the relationship between *h* and s did vary across some groupings. Specifically, the decay rate was significantly larger for catalytic genes than for any of the other categories, again indicating that mutations in these genes tend to be more recessive than those in other genes. Genes that were more highly expressed and those that tended to be more connected had a smaller decay parameter, indicating that mutations in these genes tended to be more additive (Fig. 3C and 3D). Strikingly, we could not reject a model where structural genes had the same decay parameter as highly connected genes and non-structural genes that are both highly connected and have high levels of expression (Fig. 3D). These results argue that structural genes do not appear to have a unique *h-s* relationship. Rather they share the properties of other genes that are both highly connected and have a high level of expression.

Our results motivate further development of a more general model for dominance. We extended the model of Hurst and Randerson (9). In our model, fitness, *f(x)*, for a given level of gene expression *x*, is described by:

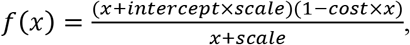

where the *intercept* relates to the functional importance of a given gene and together with the *scale*, determines the optimal expression level of the gene. We assume that gene expression comes at a fixed *cost* per unit expression level. We compute *s* and *h* from this model based on how reducing the expression level by one half (for heterozygotes) or completely (for the homozygotes) affects fitness (18) (fig. S11). Under this model, a non-essential gene where few molecules are needed for optimal function (solid blue curve in Fig. 4A) will have a wild-type fitness at a low expression level (solid point). Reducing the amount of active protein by one half (the assumed impact of a deleterious heterozygous mutation) will only slightly decrease fitness, resulting in a recessive mutation. In contrast, for an essential gene where many molecules are needed, the fitness function will be much flatter and the optimal expression will be much higher (dashed yellow curve in Fig. 4A). Here, reducing the amount of active protein by one half will result in a larger decrease in fitness, implying that mutations will be more additive. Simulations under our model (*18*) recapitulate the key features seen in our empirical data (Fig. 4B and Fig. 4C). Specifically, while all genes are predicted to show a *h-s* relationship under our model (fig. S12), this relationship will be less-steep in genes with a higher optimal expression level (orange points in Fig 4B), indicating that for a given selection coefficient, genes with high expression will tend to be more additive. Stratifying by realized gene expression level shows qualitatively similar patterns—mutations in genes with higher expression levels are predicted to be more additive (Fig. 4C).

**Fig 4.**
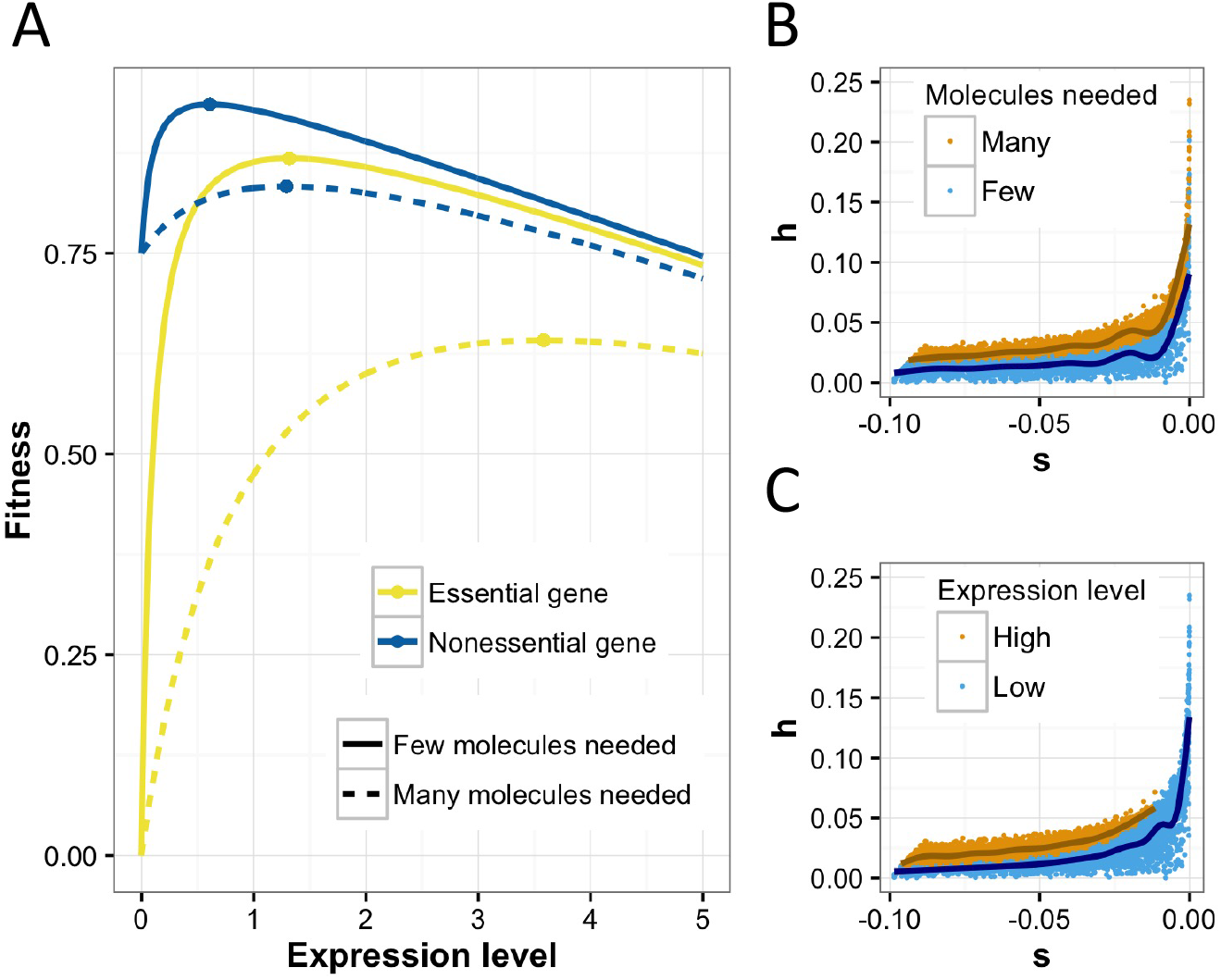
A new, comprehensive model for the evolution of dominance. **(A)** The relationship between fitness and expression level (arbitrary units). Note, a fitness cost for increasing gene expression is assumed (see 18). **(B)** Predicted *h-s* relationship when many molecules (orange) and few molecules (blue) are needed. **(C)** Predicted *h-s* relationship when the expression level is high (orange) and low (blue). Note that the patterns predicted in (**B** and **C**) mirror those seen empirically in our analysis.

Overall, our work provides a fine-scale molecular population genetic demonstration using genetic variation data from natural populations that more deleterious mutations tend to be more recessive than less deleterious mutations. Further, while finding some support for the popular metabolic network theory of dominance, we find that it is insufficient to explain patterns of dominance in all types of genes. Rather, our results support a more general model for the occurrence of dominance. Specifically, our findings suggest that dominance and the *h-s* relationship arose as a natural and inevitable outcome of the functional importance of genes and their optimal expression level. In addition, under our model, dominance can evolve in haploid organisms, passing a previous test of the evolution of dominance that rejected both Fisher's and Haldane's original models (20).

Our findings have implications for evolutionary and medical genetic studies. First, many deleterious mutations tend to be recessive, and may accumulate in heterozygotes and be maintained in populations, which could increase the role of population history in affecting patterns of deleterious mutations and the genetic load (1, 2). Second, the location of a gene in a biological network and optimal expression level will influence both the selection coefficient and prone to having fitness effects and being potentially involved in complex traits, consistent with the recently proposed omnigenic model (21).

## Acknowledgements

The data used in this paper is archived at the European Nucleotide Archive study accession numbers PRJEB19780 and PRJNA284572. The code used for analysis is available at www.github.com/LohmuellerLab/dominance. We thank Dan Balick for detailed comments on the manuscript. This work was supported by a Searle Scholars Fellowship and NIH Grant R35GM119856 (to K.E.L.).

## Supplementary Materials

### Materials and Methods

#### Data

We collected sequencing data for 13 *A. lyrata* plants from Novikova et al 2016 (16) and sequencing data for 16 *A. thaliana* plants from Durvasula et al 2017 (17). We aligned accessions to their respective genomes *(A. thaliana* to TAIR10 (22) and *A. lyrata* to the JGI reference sequence v1.0 (23)) using BWA-MEM (BWA 0.7.7-r441) (24) with a penalty of 15 for unpaired read pairs. We removed duplicated reads using Picard v2.7 and performed local indel realignment using Genome Analysis Toolkit (GATK v3.6) IndelRealigner (25). We called SNPs using UnifiedGenotyper and filtered variants using the recommendations from GATK:

QualByDepth < 2.0 || FisherStrand > 60.0 || RMSMappingQuality < 40.0 || MappingQualityRankSumTest < -12.5 || ReadPosRankSum < -8.0 || StrandOddsRatio > 3.0 || HaplotypeScore > 13.0

We annotated SNPs using SnpEff v4.3a (26). We used gene annotations (TAIR10) to filter only coding sequences (CDS) and created site frequency spectra (SFS) for synonymous and nonsynonymous variants separately. We calculated folded SFSs in order to avoid assigning an ancestral allele, which is difficult to do in these species due to extensive genome rearrangements (23).We downsampled the SFS in *A. lyrata* from 13 entries to 11 using a hypergeometric downsampling scheme (27).

We ensured that population structure did not affect our frequency spectra by performing principal components analysis (PCA) and checking the distribution of pairwise differences between samples. We removed samples that were highly related within each species as determined by outliers in the number of pairwise differences and individuals that cluster very closely on the PCA run on the genotypes (28) (Fig. S3). When two accessions were closely related, we retained one individual selected at random. For the *A. thaliana* dataset, we removed samples 35601, 35513, 35600, 37469 and for the *A. lyrata* dataset, we removed samples SRR2040788, SRR2040795, SRR2040829.

We annotated each coding site according to the gene name and gene ontology (GO) term and subset the data into different GO term categories to perform our inference of dominance and the DFE separately on these categories. We annotated each gene based on connectivity and gene expression. Connectivity was determined by the STRING database v10 (29). We downloaded the *A. thaliana* (organism 3702) protein network data and restricted our analysis to high confidence (>0.7) interactions. Connectivity is then equally subdivided into three categories: low connectivity, intermediate connectivity, and high connectivity (e.g. Fig. 3). We obtained expression data for *A. thaliana* from the 1001 Epigenomes project (NCBI GEO: GSE80744; (30)), which provides a processed read count matrix for each gene across all accessions. We obtained the median expression value across all accessions, and arrived at a single value for each gene. Expression level is then equally subdivided into three categories: low expression, intermediate expression, and high expression (e.g. Fig. 3).

#### Models of dominance and likelihood ratio test

We test three different models of the relationship between the selection coefficient of a mutation (*s*) and the dominance coefficient (*h*). Here, s and *h* are defined such that the fitness of the homozygous wild-type genotype is 1, the fitness of the heterozygous genotype is 1+*hs*, and the fitness of the homozygous mutant genotype is 1+s. The first model assumes that *h* is 0.5 and does not depend on s (additive model). The second model assumes that *h* is independent of s, but different from 0.5 (constant *h* model). This model allows for dominant or recessive mutations. The third model assumes a functional relationship between *h* and *s* (*h-s* relationship model). We model this relationship with two parameters according to the following equation:

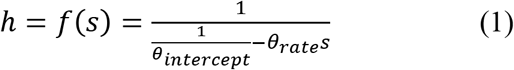

The first parameter, *θ*_intercept_, defines the value of *h* at *s* = 0. The second parameter, *θ*_rate_, defines how quickly *h* approaches zero with decreasing negative selection coefficient (see Fig. 1D). We assume that *θ*_rate_ is positive. Large positive values of *θ*_rate_ imply that *f*(s) quickly approaches *h*=0, and even slightly deleterious mutations are recessive. Small positive values of *θ*_rate_ imply that only strongly deleterious mutations are recessive.

Overall, we assume that the DFE of new mutations (i.e. the distribution of s) follows a gamma distribution (31-33). Thus, the additive model has two DFE parameters (shape and scale of the gamma DFE) and no dominance parameters, since we fix *h* to be 0.5. The constant *h* model has one additional parameter, the value of h. The *h-s* relationship model has two additional parameters, *θ*_intercept_ and *θ*_rate_. Note that when *θ*_rate_ approaches zero, the *h-s* relationship model of eq. 1 converges to the constant *h* model, and when *θ*_rate_ approaches zero and *θ*_intercept_ approaches 0.5, the model converges to the additive model. Thus, the three models are nested, and we can formulate a likelihood ratio test based on maximum log likelihoods (LL) comparing the three different dominance models. The test statistic Λ is defined as 2(LL_H1_ - LL_H0_), where H0 is the null hypothesis (either additivity or constant *h*) and H1 is the alternative hypothesis (either constant *h* or *h-s* relationship). The statistic Λ is asymptotically chi-square distributed, with degrees of freedom equal to the difference in the number of parameters between the null and the alternative model. Thus, we formulate three different tests:

1. Testing the constant *h* model (H1) against the additive model (H0).
2. Testing the *h-s* relationship model (H1) against the additive model (H0).
3. Testing the *h-s* relationship model (H1) against the constant *h* model (H0).

#### Population genetic inference of dominance using data from a single outcrossing population

We developed a Poisson random-field model of polymorphisms (14) for estimating the parameters in the models described above. We assume that nonsynonymous mutations are under the effects of purifying selection, and we assume that synonymous mutations are neutral. We present two approaches to estimate these parameters from the data: 1) estimating dominance using data from a single outcrossing population (e.g. *A. lyrata)*, and 2) using data from both an outcrossing (e.g. *A. lyrata)* and a highly inbreeding population (e.g. *A. thaliana*) simultaneously to estimate dominance. We start by presenting the first approach.

To account for the effects of changes in population size on the nonsynonymous SFS that might confound estimates of selection, we first estimate a demographic model using the synonymous SFS (34). Selection parameters are then estimated conditional on the estimated demographic model. Previous work has shown that this approach leads to unbiased estimates of the selection parameters by controlling for background selection, selective sweeps, and hidden population structure (32, 35).

In short, we infer the parameters of a population size change model using the synonymous site frequency spectrum (SFS) under the Poisson Random Field framework (see Huber et al. (32) and Kim et al. (35) for details). For both species that we analyzed *(A. lyrata* and *A. thaliana)*, a three-epoch model with three discrete size changes fits better to the synonymous SFS than a two-epoch model or a constant population size model (Table S1, Fig. S1). Thus, all subsequent inferences use the three-epoch model.

Conditional on the estimated demographic parameters of the three-epoch model, we next use the nonsynonymous SFS to estimate the selection parameters, i.e. the shape and scale parameter of a gamma distributed DFE (Θ_DFE_), and the rate and intercept parameter of the *h-s* relationship, Θ_h_ = {*θ*_intercept_, *θ*_rate_}. We use the Poisson likelihood to estimate the combined vector of parameters {Θ_DFE_, Θ_h_}. The likelihood is calculated as

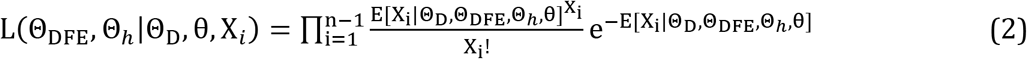

Here, Θ_D_ is a vector of demographic parameters, X_i_ is the count of SNPs with frequency *i* in the sample (the entries of the SFS), *θ* is the population mutation rate, and *n* is the sample size.

We set Θ_D_ to the maximum likelihood estimates of the demographic parameters Θ_D_, and *θ* to the nonsynonymous population scaled mutation rate, *θ*_NS_ = 4N_e_ μL_NS_. We estimated *θ*_NS_ from *θ*_s_ by accounting for the difference in synonymous and nonsynonymous sequence length.

The expected values of X_i_ refer to the expected entries of the SFS given demography and selection parameters. We used the software ∂a∂i (27) to compute the expected SFS for a 2dimensional grid of 1 million pairs of Nes and *h* values on grid that is exponential in Nes and linear in *h* (see also *Cubic spline interpolation to speed up the computation of cached SFS)*. We vary *h* from zero (completely recessive) to one (completely dominant), and N_e_s from -N_e_ (i.e. lethal) to -1x10^-4^ (effectively neutral). This set of site frequency spectra is then used to calculate the expected SFS for an arbitrary distribution of N_e_s and *h* values. This is done by numerically integrating over the respective spectra weighted by the gamma distribution. Since we assume one N_e_s value corresponds to a single *h* value (equation 1), this is a one-dimensional integration. The numerical integration was done using the ‘numpy.trapz, function as implemented in ∂a∂i.

Numerical optimization is used to find the parameters of the DFE and dominance model that maximize the Poisson likelihood (equation 2). For this optimization step, we use the BFGS algorithm as implemented in the ‘optimize.fmin_bfgs, function of scipy. To avoid finding local optima, we repeated every estimation approach from 1000 uniformly distributed random starting parameters. Our approach allows us to estimate the parameters of any arbitrary distribution of N_e_s values and any arbitrary function that relates *h* to s (or *N_e_s)*.

To summarize, our inference of dominance and DFE parameters (Θh, ΘDFE) consists of the following steps:

1. Infer the parameters of a demographic model and the effective (ancestral) population size for the outcrossing population.
2. Conditional on the demographic model, compute the expected SFS for a 2D grid of *h* and *N_e_s* values.
3. Start at a certain vector of dominance and DFE parameters (Θ_h_, Θ_DFE_). Note that the DFE here is defined in units of s, not *N_e_s*.
4. Compute the DFE in units of *N_e_s* by scaling the DFE from step 3 by the respective ancestral population size.
5. Compute the *h* value for the grid of *Nes* values according to eq. 1 and the parameters Θ_h_. Then use the 2D lookup table generated in step 2 to find the closest SFS for each pair of *h* and *N_e_s*. Integrate those SFS after weighting according to the DFE to find the expected SFS given the DFE and *h-s* relationship.
6. Given the expected and the empirical SFS for the outcrossing population, compute the log likelihood according to eq. 2.
7. By repeating steps 3-6, the log likelihood can be calculated for an arbitrary set of parameters. Maximum likelihood parameters are computed numerically by maximizing the likelihood using iterative non-linear optimization methods such as BFGS or Nelder-Mead (36).

The ancestral effective population size in step 4 is calculated from the demographic model. Fitting the demographic model to the synonymous SFS provided an estimate of *θ*_S_ = 4N_e_μL_S_ for synonymous sites, where μ is the neutral per base-pair mutation rate and L_S_ is the synonymous sequence length. Using this formula, we estimated *N_e_* by setting the neutral mutation rate to 7 x 10^-9^ (37). Note that when partitioning our data into different gene categories and estimating the selection parameters for each category separately, we also allow for a different ancestral *Ne* and demographic estimates in those categories to control for different levels of background selection in different genomic regions (38-41).

Finally, we can compute the likelihood at the maximum likelihood parameter values for the three different dominance models (i.e. additive model, constant *h* model, and *h-s* relationship model), and compute the likelihood ratio test statistic Λ, which will allow for model comparison.

#### Cubic spline interpolation to speed up the computation of cached SFS

Step 2 in our inference method involves computing a lookup table of one million SFS for a wide range of 1000x1000 pairs of *N_e_s* and *h* values. Although each single computation of a SFS is relatively fast, it is computationally expensive to compute the total of one million SFS with ∂a∂i. We sped up this computation by utilizing the fact that the SFS across close *N_e_s* and *h* values is fairly smooth. Thus, we only compute the expected SFS for a coarse grid of 50 x 20 *N_e_s* and *h* values, and then interpolate the entries of the SFS for a much finer grid of 1000 x 1000 *N_e_s* and *h* values. The interpolation is done using the CubicSpline function of the python package scipy.interpolate. Each frequency of the SFS is interpolated separately in a two-step process: first, each frequency is interpolated for 1000 positions along the *Nes* axis, keeping *h* constant, leading to a grid of 1000 x 20 SFS. Then, each frequency is interpolated along the *h* axis, keeping *N_e_s* constant, leading to the final grid of 1000 x 1000 SFS. Examples of the cubic spline interpolation of frequency classes of the SFS along the *N_e_s* and *h* axes demonstrate that the interpolation works well for a wide range of h, *N_e_s* and minor allele frequency (MAF) values (Fig. S5 and S6).

#### Population genetic inference of dominance using data from an outcrossing and a highly inbred population

The nonsynonymous SFS for different values of *h* can be very similar when modifying the selection coefficient accordingly (see Fig. 1A). This suggests that the power for estimating dominance might be small when using only data from a single outcrossing population. This can be seen in Fig. S2A, where simulations with h=0.5 (H0) are compared to simulations with a constant *h* of 0.46 (H1). Such a small difference in *h* leads to a considerable overlap in the distribution of the likelihood ratio test statistic Λ between simulations under H0 and H1, and there is no power to discriminate those two hypotheses.

We propose to increase power for detecting the true dominance model, and improve parameter estimation, by combining data from an outcrossing species with data from a selfing species. The main factor determining the SFS of the outcrossing species is the difference in fitness between the homozygous wild-type and the heterozygous genotype, having fitnesses 1 and 1-hs, respectively. The difference in fitnesses between these two genotypes affects the SFS because deleterious mutations are segregating at low frequencies and thus random mating rarely produces homozygous derived genotypes. On the other hand, for a strongly selfing species, genotypes mostly are in the homozygous state due to the high level of inbreeding. The main factor determining the SFS in the selfing species is the difference in fitness between the two homozygous genotypes, having fitnesses 1 and 1-s, respectively. Thus, the data from the outcrossing species mainly provides information about the product of *h* and s, whereas the data from the selfing species provides information about s independent of h. Combining information from both datasets therefore should allow us to estimate dominance with higher accuracy than either species alone.

To extend our inference to an inbreeding/outcrossing pair of populations, we need to calculate the likelihood of the parameters given the nonsynonymous SFS of both populations. When the two species are strongly diverged such that they do not share ancestral polymorphisms, the allele frequencies are independent and the likelihood can be computed as the product of the probability of the outcrossing SFS (SFS_o_) and the probability of the inbreeding SFS (SFS_I_). In terms of log-likelihoods (LL), this equates to:

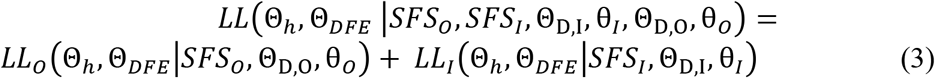

The first term of the sum, the log likelihood of the selection parameters (Θ_h_ and *Θ_DFE_)* given the outcrossing SFS, is computed using the approach developed above for the case of a single outcrossing population. To calculate the log likelihood for the inbreeding SFS (the second term of the right hand side of equation 3), we need to account for the effect of inbreeding on the SFS. For strongly inbred species such as *A. thaliana* with a selfing rate of at least 97% (42), we assume that the inbreeding coefficient *F* is effectively 1. In this case, the diffusion equation model reduces to a scaled additive model. This can be derived from the formulas of the mean and variance of the change in frequency at an allele frequency p: *M(p)* and *V(p)*. In the most general case, with arbitrary inbreeding and dominance, these two quantities are (43):

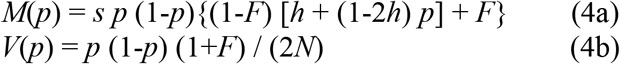

In the case of additive mutations in an outcrossing population (*F*=0, *h*=0.5), these quantities become:

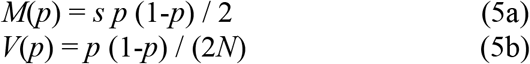

In the case of a highly inbred population with arbitrary dominance (F=1) these quantities become independent of *h*:

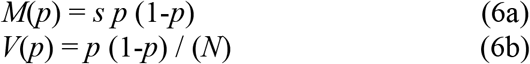

The equations for the case of *F*=1 (eq. 6a,b) is just a scaled version of the equations for additive mutations in an outcrossing population (eq. 5a,b), with twice the change in mean allele frequency (eq. 6a), and twice as much drift (eq. 6b). This allows us to use the framework of ∂a∂i, developed for outcrossing populations, and apply it to data from highly selfing populations.

We need to take into account the effect of inbreeding on *M(p)* and *V(p)* according to eqs. 6ab. The effective population size that we estimate with ∂a∂i based on the synonymous SFS is already taking into account the effect of inbreeding on *V(p)*, since it is the population size that effectively generates the same amount of drift as the standard Wright-Fisher outcrossing model assumed by ∂a∂i (i.e. eq. 5b). Next, we multiply s by a factor of 2 to find the effective selection coefficient *s_e_*. Finally, we use these effective parameters, *s_e_* and *N_e_* to compute the expected SFS for the highly selfing population using the framework of ∂a∂i.

The full inference of a common set of dominance and DFE parameters (Θ¾ *Θ_DFE_)* is similar to the steps outlined above for a single outcrossing population.

1. Infer the parameters of a demographic model and the effective (ancestral) population size for both the inbreeding and the outcrossing populations. This is done independently for the two populations.
2. Conditional on the demographic model of the outcrossing population, compute the expected SFS for a 2D grid of *h* and *N_e_s* values. For the inbreeding population, compute the expected SFS for a 1D grid of *N_e_s* values, fixing *h* to 0.5.
3. Start at a certain vector of dominance and DFE parameters (Θ¾ *Θ_DFE_)*. Note that the DFE here is defined in units of s, not *N_e_s*.
4. Compute the DFE in units of *N_e_s* by scaling the DFE with the respective population size separately for the inbreeding and the outcrossing population. For a gamma distributed DFE, this amounts in multiplying the scale parameter by *N_e_*.
5. For the inbreeding population, additionally scale the DFE from step 3 by a factor of 2 to derive the effective DFE in units of *N_e_s_e_*.
6. For the outcrossing population, compute the *h* value for the grid of *N_e_s* values according to eq. 1 and the parameters Θ*h*. Then use the 2D lookup table generated in step 2 to find the closest SFS for each pair of *h* and *N_e_s*. Integrate those SFS after weighting according to the DFE to find the expected SFS given the DFE and *h-s* relationship.
7. Compute the expected SFS for the inbreeding population by integrating across the 1D lookup table of SFS after weighting each SFS according to the DFE in units of *N_e_s_e_*. Note that *h* is fixed to 0.5.
8. Given the expected and the empirical SFS for both the inbreeding and the outcrossing populations, compute the log likelihood according to eqs. 2 and 3.
9. By repeating steps 4-8, the log likelihood can be calculated for an arbitrary set of parameters. Maximum likelihood parameters are computed numerically by maximizing the likelihood using iterative non-linear optimization methods such as BFGS or Nelder-Mead (36).

#### Bootstrapping and testing model parameters

Our maximum likelihood approach of inferring the DFE and dominance parameters only returns a point estimate, and does not include a measure of the uncertainty of the estimate. Further, since the approach numerically optimizes the likelihood and estimates demographic parameters, numerical errors might lead to a larger uncertainty in parameters than expected based on the shape of the likelihood function. Thus, we follow a non-parametric bootstrapping approach by Poisson resampling both the synonymous and nonsynonymous empirical SFS and re-estimating the demographic and selection parameters for each resampling. From 20 bootstrapped parameters we then compute the standard error and the 95% confidence interval (Fig. 3C). To test for difference in certain parameters between gene categories, we computed a z-score by dividing the difference in the estimate by the estimated standard error of the difference. The *P*-value is then computed based on the standard normal distribution (Fig. 3D).

#### Robustness in establishing the *h-s* relationship

The negative relationship between *h* and *s*, such that more deleterious mutations are more recessive, was first reported by a series of mutation accumulation (MA) experiments in *Drosophila* (10, 11), and later supported by two studies in yeast (13, 19). However, the validity of the results was questioned (19, 44). Further, the more comprehensive and detailed study in yeast restricts their *h-s* relationship models such that more deleterious mutations are only allowed to become more recessive than less deleterious mutations, but not more dominant (*19*). Such a study, by definition, cannot find support for a positive relationship between *h* and s, because the model did not allow for such a relationship. Thus, based on previous work, it has not clearly been established that more deleterious mutations become more recessive.

Therefore, we also tested an alternative model where *h* converges to one instead of zero, i.e. more deleterious mutations are more dominant than less deleterious mutations:

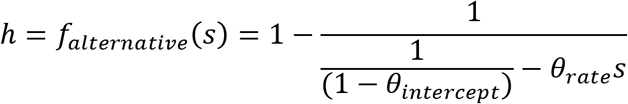

However, this model does not improve fit to the SFS over a constant *h* model or the *h-s* relationship model of equation 1. When using only data from *A. lyrata*, then the log likelihood of the alternative *h-s* relationship model (LL = -405.1) is similar to that of the constant *h* model (LL = -404.9), and much lower than the log likelihood of the *h-s* relationship model of equation 1 (LL = -218.8). The small estimated *θ_rate_* parameter (3,980) suggests that this model is equivalent to the constant *h* model where *h* does not change with *s*. Similar results are obtained with our two-population inference, using data from both *A. lyrata* and *A. thaliana*. Again, the log likelihood of the alternative *h-s* relationship model (LL = -885.2) is similar to that of the constant *h* model (LL = -885.3), and much lower than the log likelihood of the *h-s* relationship model of equation 1 (LL = -399.7). The extremely small estimated *θ_rate_* parameter (0.26) suggests that this model is equivalent to the constant *h* model. Thus, in summary, we conclude that a model where more deleterious mutations become more dominant does not fit the SFS as well as a model where more deleterious mutations become more recessive.

#### Robustness of inference to model mis-specifications

When we make simultaneous use of data from both outcrossing *(A. lyrata)* and inbreeding *(A. thaliana)* species for inferring dominance, we implicitly make the assumption that the DFE is the same in both species. However, for highly diverged species such as humans and *Drosophila*, it was shown recently that the DFE, in units of s, is significantly different (32). One potential concern is that differences in the DFE between species could lead to falsely inferring an *h-s* relationship when the true model is additivity.

However, we found additional support for the of *h-s* relationship model. First, we see significant support for an *h-s* relationship over an additive or constant *h* model even when basing our inference only on the outcrossing *A. lyrata* data (Table S2). Further, the estimates of the DFE and dominance parameter estimates agree reasonably with each other across different ways of doing the inference. Specifically, estimates made using only *A. lyrata*, agree with those using *A. lyrata* and *A. thaliana* combined, although in the former case, the confidence limits are wider (Fig. S8). Second, we explored the effect of different DFEs on our inference procedure using simulations. We ran simulations under an additive model, with parameters of the DFE taken from separate estimates of the DFE in each species (for details see the next section). Then, on each simulated dataset, we fit the demographic and selective models. Lastly, we compute the sum of log likelihoods *(LL_O_ +LL_I_)* assuming a unique additive DFE in both species (true model). Then we compare this log likelihood to the log likelihood that assumes the same DFE, but an *h-s* relationship (incorrect model). We find that the additive log likelihood always sums up to a larger value than the log likelihood assuming the same DFE, but an *h-s* relationship. This pattern in the simulations contrasts with what is seen in the actual empirical data. For the empirical data, we find that the log likelihood of the additive model with unique DFEs *(LL_O_ +LL_I_* = -466 - 84 = - 550) is smaller (i.e. a worse fit) than the log likelihood assuming the same DFE, but an *h-s* relationship (LL = -400, see Table S3). This suggests that the additive model has a worse fit than an *h-s* relationship model, even when the assumption of an identical DFE in both species is relaxed. In summary, analyses of simulated data suggest that it is possible to distinguish between different DFEs between species and a true *h-s* relationship. It is unlikely for our inference framework to infer a spurious *h-s* relationship due to differences in the DFE between species.

Another assumption of our approach is that the inbreeding coefficient *F* of the selfing population equals 1. We tested robustness to this assumption by simulating SFS data for a selfing population with selfing rate at the lower end of what has been estimated for *A. thaliana* (97%; (42, 45-47)). We then compared this SFS to an SFS that is simulated under full selfing (F=1), and found that the SFS match up well. Similar results are found for even lower selfing rates of 90% or 85% (Fig. S9). Moreover, we found that our approach leads to unbiased estimates when simulating data under a selfing rate of 97% (Fig. S10). Thus, an inbreeding rate of 97% is high enough to ensure unbiased estimation of dominance parameters with our approach.

#### Simulation setup

To test our inference procedure, we simulated data using the forward simulation software PReFerSim (48), but changed the source code of the software to allow for an *h-s* relationship according to eq. 1. We simulate genome-wide data under the three-epoch model, with

*θ*_synonymous,Inbreeding_=41,800, *θ*_Nonsynonymous,lnbreeding_=96,600, *θ*_Synonymous,Outcrossing_=131,600, and *θ*_Nonsynonymous,outcrossing_=304,000. Here, *θ* is *4NeμL*, where *L* is the respective synonymous or nonsynonymous sequence length, μ is the neutral mutation rate, and *Ne* is the ancestral population size. Further, we simulated smaller sets of data that reflect the relatively small number of structural genes, with all values of *θ* being 10 times smaller. The simulation parameters for the DFE, the demographic model, and the *h-s* relationship are taken from the empirical estimates from the genome-wide data (see Table S1 and S3). However, the simulations are downscaled to a 50-fold smaller population size than estimated to increase the speed of the simulations (32). After simulating the respective synonymous and nonsynonymous SFS under both inbreeding and outcrossing, we estimate the demographic parameters, the DFE parameters, and the dominance parameters using our method.

We simulated 100 replicates of the following scenarios: First, we simulated under the additive model, assuming the same DFE in both populations. After running the inference, this leads to the null distribution of the test statistic Λ (Fig. 2C and 2D). Second, we simulated under the constant *h* model. This leads to the distribution of Λ under the alternative hypothesis of constant *h* (Fig. 2C). Finally, we simulated under the *h-s* relationship model. This leads to the distribution of Λ under the alternative hypothesis of an *h-s* relationship (Fig. 2D). We find that we can estimate the true parameters of the *h-s* relationship under all simulation scenarios (Fig. S10).

The two simulated null distributions of Λ in Fig. 2C and 2D follow closely to the expectation under the asymptotic theory, with only a slightly larger mean and standard deviation: the expected mean and standard deviation of a chi-square distribution with df=1 is 1 and 1.9, the observed mean and standard deviation of Λ in Fig. 2C is 1.9 and 2.9. The expected mean and standard deviation of a chi-square distribution with df=2 is 2 and 2, the observed mean and standard deviation of Λ in Fig. 2D is 2.1 and 4.9.

#### Model for how the evolution of optimal gene expression explains dominance patterns

Our model of how optimal gene expression leads to dominance is an extension of the model of Hurst and Randerson (9). In this model, dominance is a direct consequence of optimized gene expression. Fitness is modeled as a function of gene expression. Higher gene expression leads to higher fitness, but the gain from increasing gene expression is lower for higher levels of gene expression than for lower levels of gene expression (diminishing returns function). For enzymatic genes, this relationship was shown to be a consequence of metabolic pathway dynamics, assuming that the output of the system (flux) is directly related to fitness (7). For genes encoding structural proteins, it is imaginable that after enough protein is produced to build certain structures in the cell or the extracellular matrix, additional protein does not improve its functional role any further.

To formalize such a type of diminishing returns function, Hurst and Randerson assume a simple functional relationship between expression level and fitness, *f(x)* = *x*/(1+*x*), where *x* is the expression level (arbitrary units), and *f* is the fitness. Further, they assume that per unit of *x*, there is a cost associated with gene expression. In biological systems, these costs could be related to spending cellular resources (amino acids and nucleotides), allocation of cellular machineries (RNA polymerase and ribosome), or energy consumption (49). The expression cost is included as a parameter that quantifies the reduction in fitness per unit of gene expression, such that *f(x)* = *x*/(1+*x*)(1-cost\**x*). For simplicity, we assume that the cost per unit of gene expression is the same for every gene.

We now extend this model in two ways. First, the Hurst and Randerson model assumes that the fitness at zero expression level is zero. However, experiments in bacteria, yeast and a number of other organisms have shown that a considerable proportion of genes are non-essential, such that fitness would not reduce to zero when the gene is not expressed (50). We include an intercept parameter in the model that determines the fitness when the gene is not expressed. An intercept close to one indicates that the gene is non-essential and can be removed with only little reduction in fitness, whereas a value close to zero indicates that the gene is essential for survival or reproduction. Second, we add a scale parameter that allows for varying rates of increase in fitness with expression level (Fig. S11). We define the scale parameter as the expression level at which fitness is exactly in the middle between the fitness at zero expression and at infinite expression (assuming no expression costs). In biological terms, this parameter is related to the amount of protein needed by the organism to function properly. For structural proteins, many molecules might be needed to build structures in or out of the cell, which would be reflected in a large scale parameter. For enzymatic proteins, a single protein can catalyze the same chemical reaction over and over again, thus only a small amount of molecules might be needed and the scale parameter would be small. The relation between expression level and fitness is then a function of cost, intercept, and scale:

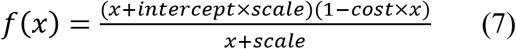

The optimal gene expression under this model can be computed by setting the derivative of *f*(*x*) to zero and solving for (positive) *x*:

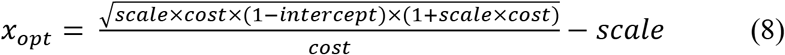

We assume that gene regulatory sequence is optimally evolved, such that genes are expressed at the level *x_opt_* (eq. 8). Next, we investigate the fitness effect of gene mutations that cause the protein to be non-functional. If the mutation is heterozygous, then the amount of functional protein is only half of the amount in the wild-type homozygous genotype. If the mutation is homozygous, then no functional protein is produced. The fitness consequences of heterozygous mutations are computed by setting gene expression x to x_opt_/2 in eq. 7. The fitness consequences of homozygous mutations are commutated by setting *x* = 0. The selection coefficient s and the dominance coefficient *h* are then defined as:

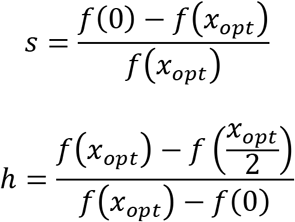

Both *s* and *h* are determined by the three parameters: *cost, intercept*, and *scale*. We can investigate the relationship between s and *h* as a function of these three parameters (Fig. S12).

Two predictions of the model can be noted: First, there is a negative relationship between *h* and *s*. More strongly deleterious mutations are more recessive than less deleterious mutations (Fig. S12). Note that this is a consequence of selection for optimal gene expression, not because of direct selection on a dominance modifier. Direct and indirect models of selection for dominance were criticized by Orr, who has noted that a predominantly haploid organism would not be able to evolve dominance (20). In at least one such organism, dominance of mutations is observed, arguing against models of selection for dominance (20). However, our model does not rely on evolution in a diploid organism, since it does not rely on selection happening only in the diploid state (see also Hurst and Randerson). It is thus in agreement with Orr’s finding. Second, although the model predicts that mutations are recessive, mutations become slightly less recessive when increasing the scale parameter, i.e. when increasing the optimal expression level of the gene. This predicts that mutations in genes with high optimal gene expression (many molecules are needed) would be more additive than genes with low optimal gene expression (few molecules are needed). This prediction matches our empirical analyses. We found that gene sets with high expression level and/or high connectivity (i.e. many molecules needed), tend to be more additive compared to gene sets having low expression level and/or low connectivity (i.e. only few molecules needed). Further, we found that mutations in genes encoding structural proteins tend to be more additive than those in genes encoding catalytic proteins (Fig. 3C).

For the simulations in Fig. 4B and 4C we simulated 5000 genes with random *intercept* and *scale* parameters and computed *h* and s of potential mutations in each gene. The cost parameter was fixed to 0.001. The *intercept* parameter was sampled from a uniform distribution with values ranging from 0.9 to 1, reflecting the fact that most new mutations are effectively neutral (35). The *scale* parameter was sampled from the absolute values of a normal distribution with mean and standard deviation of 0.1, leading to variation in the levels of optimal gene expression that is slightly skewed to lower values (i.e. assuming more genes with small optimal gene expression than with large optimal gene expression).

**Fig. S1.**
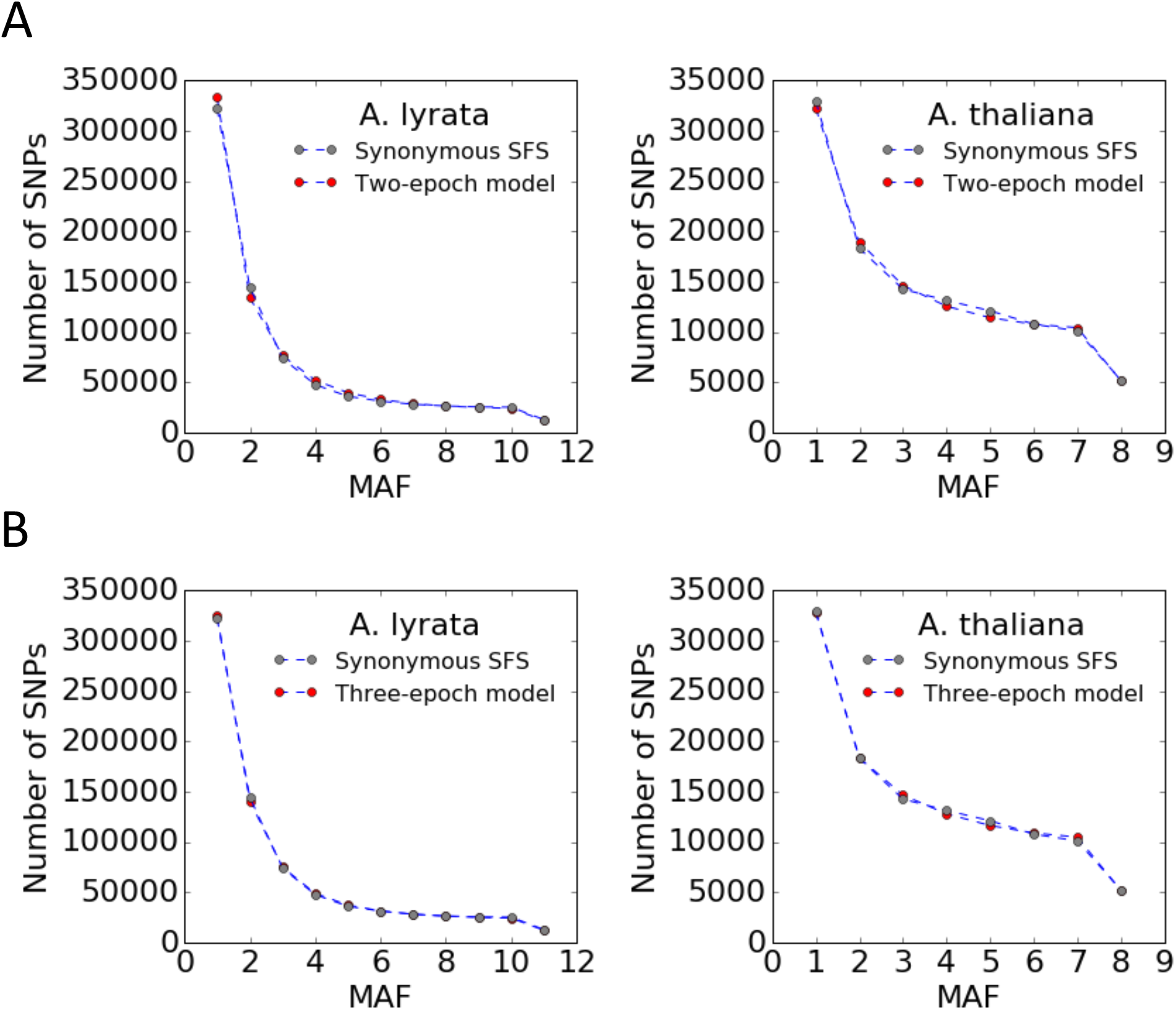
Demographic model fit to the synonymous SFS. MAF is the minor allele frequency. In both species, the three-epoch model (B) fits singletons and doubletons better than the two-epoch model (A).

**Fig. S2.**
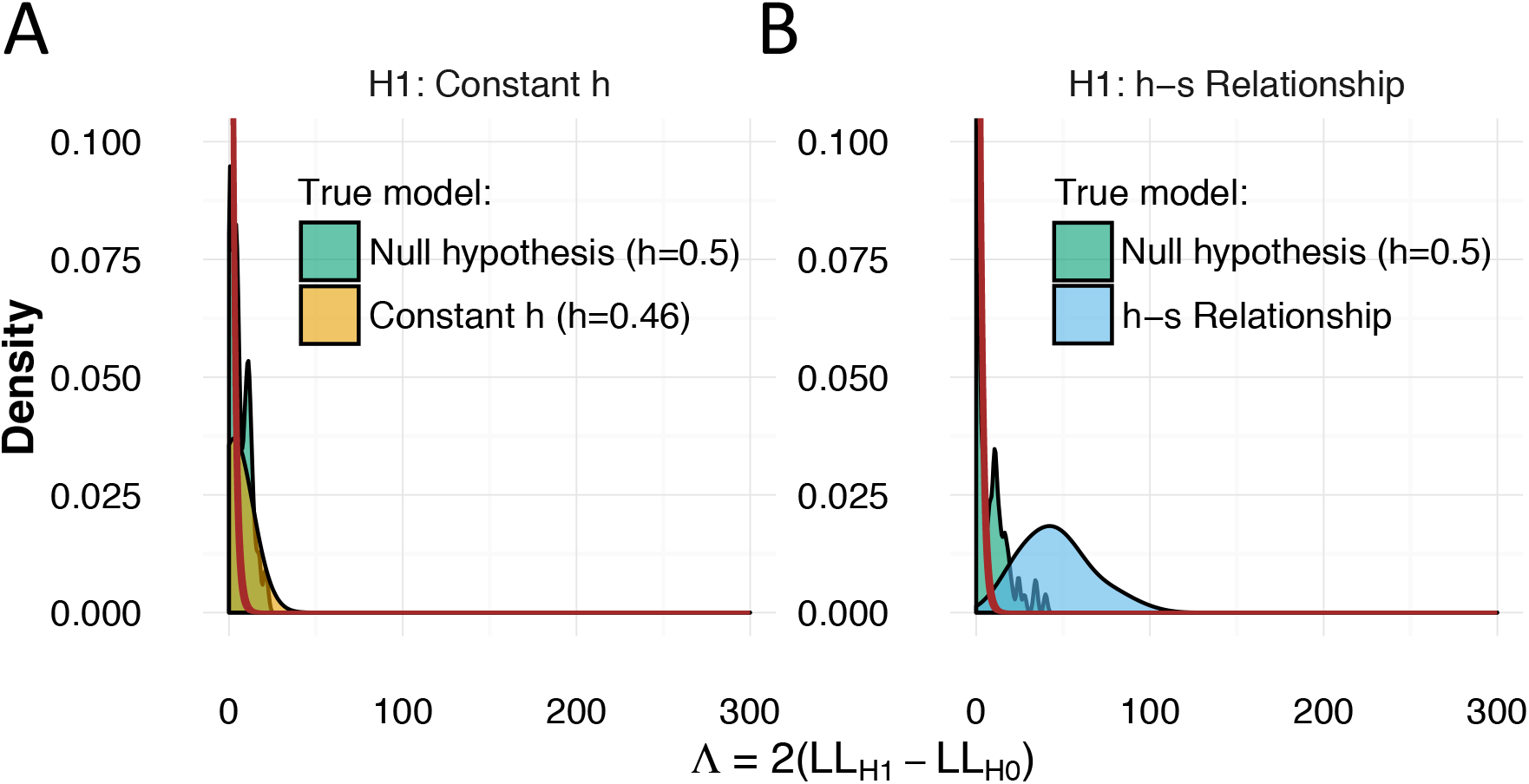
Power for discriminating between dominance models using data from a single outcrossing species *(A. lyrata)*. (A) Likelihood ratio tests comparing a constant *h* model to an additive model. When data are simulated under an additive model (green), *Λ* nearly follows a chi-square (2 *df)* distribution (red line). When the data are simulated under a model with h=0.46 (tan), the distribution of *Λ* overlaps considerably, indicating little statistical power. (B) Likelihood ratio tests comparing the *h-s* relationship model to an additive model. When data are simulated under an additive model (green), *Λ* nearly follows a chi-square (2 *df*) distribution (red line). However, when the data are simulated under the *h-s* relationship model (tan), the distribution of *Λ* is substantially larger, indicating good statistical power.

**Fig. S3.**
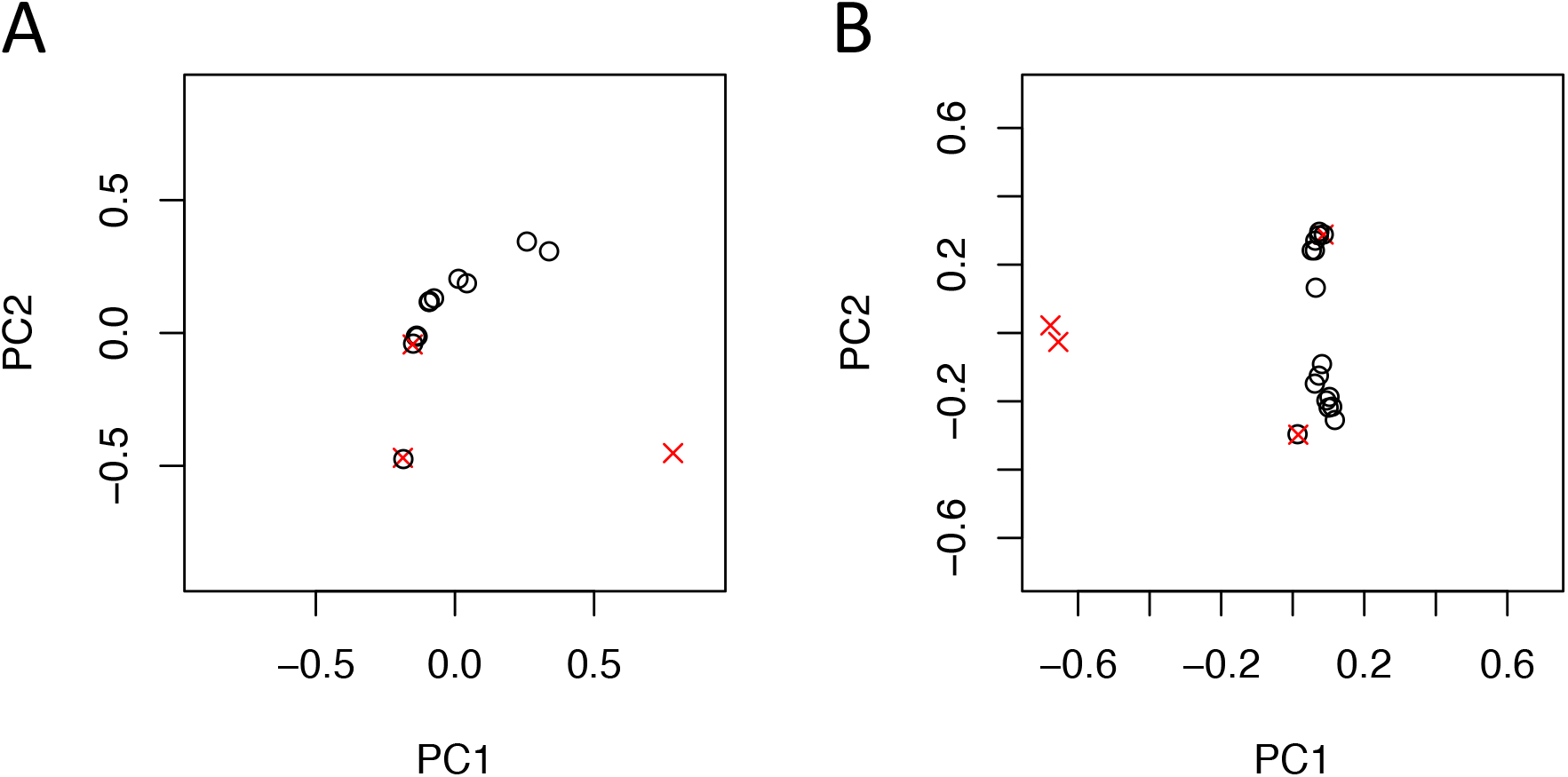
Principal component analysis of population structure. Principal component analysis (PCA) of the genetic structure of (A) *A. lyrata* and (B) *A. thaliana*. When two accessions were closely related, we retained one individual selected at random. We also removed accessions that are highly diverged from the majority of individuals. The accessions that we removed are indicated by red crosses.

**Fig. S4.**
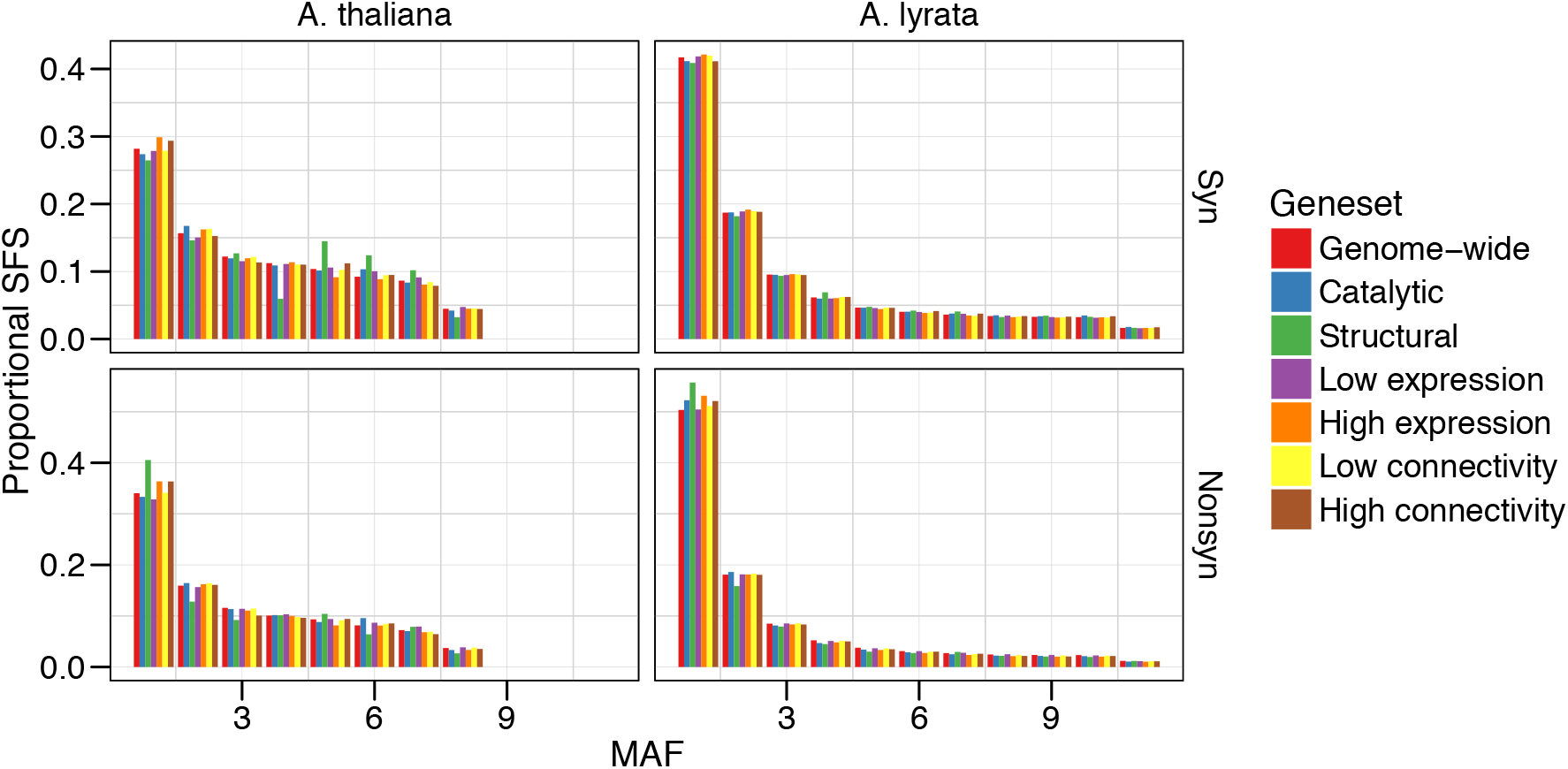
Folded site frequency spectra (SFS) for different categories of genes. In both species, structural proteins have the highest proportion of nonsynonymous singletons, suggesting these genes have experienced a greater effect of purifying selection.

**Fig. S5.**
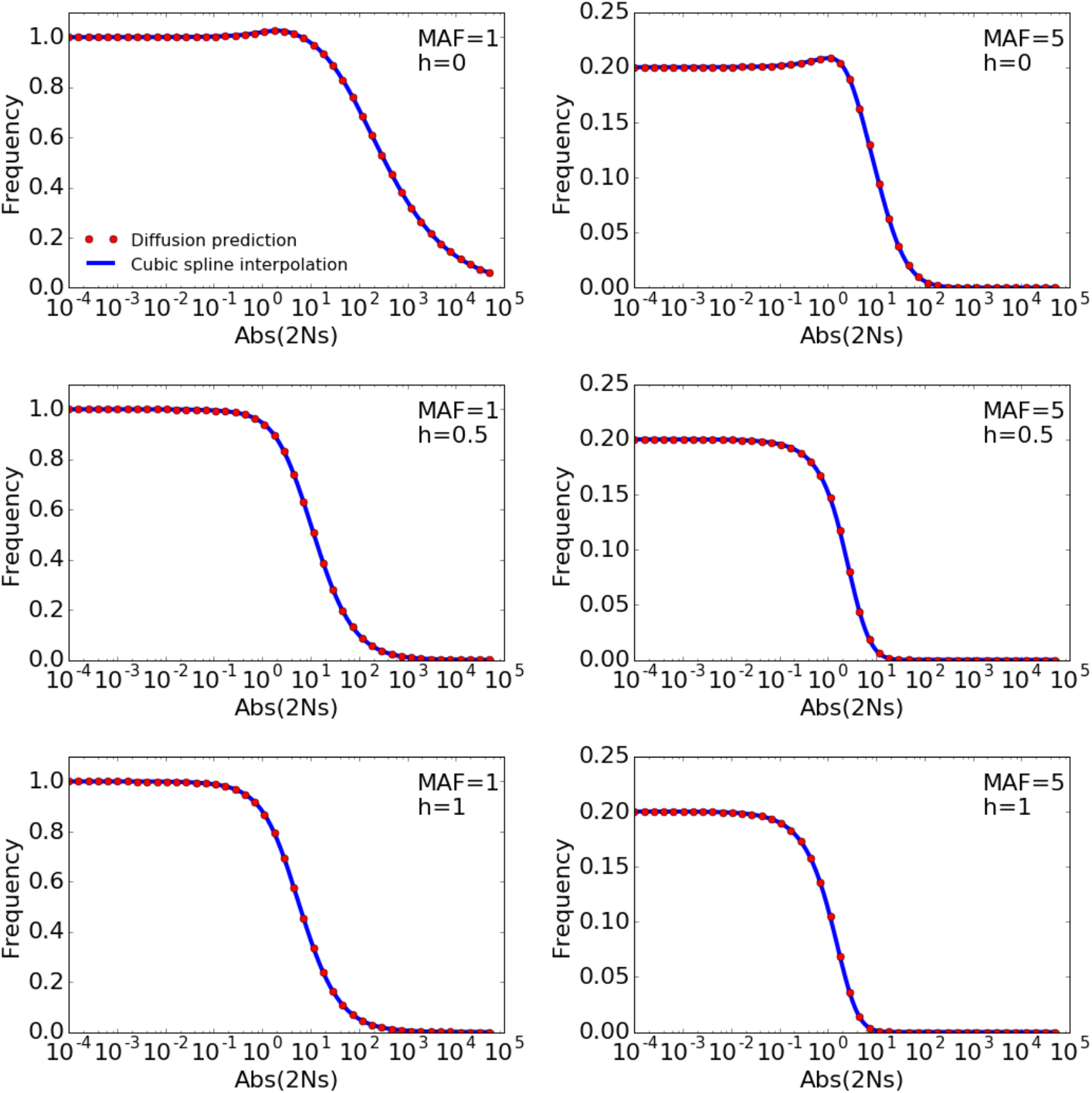
Cubic spline interpolation of the SFS along the *N_e_s* axis. Examples of cubic spline interpolation of two entries of the SFS (MAF=1 and MAF=5) for h=0, h=0.5, and h=1. The blue line is the cubic spline interpolation to the red points, which indicate the expected values under the diffusion approximation as predicted by ∂a∂i. The demography is assumed to be a constant size model. In all cases, the interpolation line fits well to the red points.

**Fig. S6.**
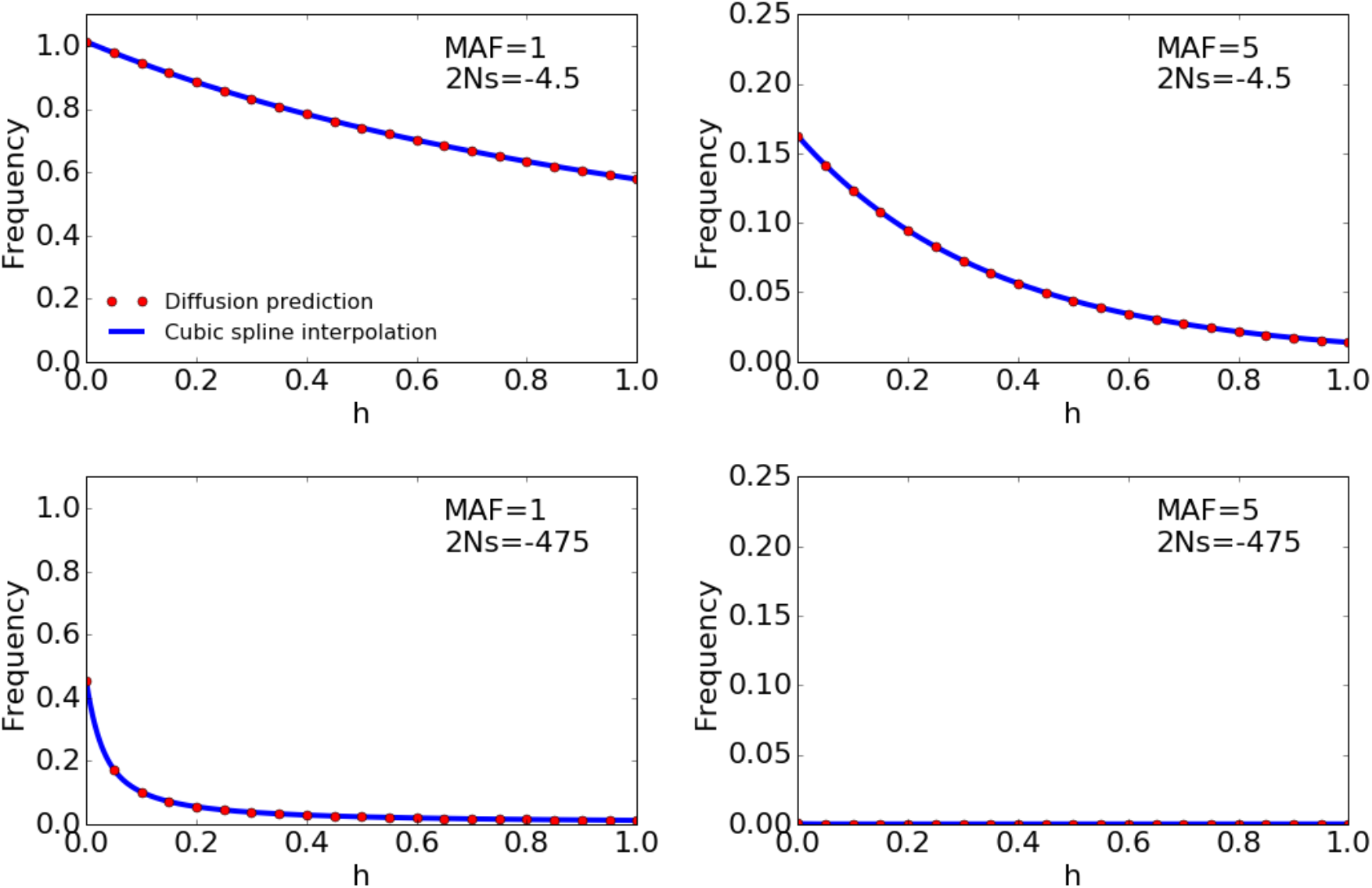
Cubic spline interpolation of the SFS along the *h* axis. Examples of cubic spline interpolation of two entries of the SFS (MAF=1 and MAF=5) for slightly deleterious (2Ns=-4.5) and strongly deleterious (2Ns=-475) mutations. The blue line is the cubic spline interpolation to the red points, which indicate the expected values under the diffusion approximation as predicted by ∂a∂i. The demography is assumed to be a constant size model. In all cases, the interpolation line fits well to the red points.

**Fig. S7.**
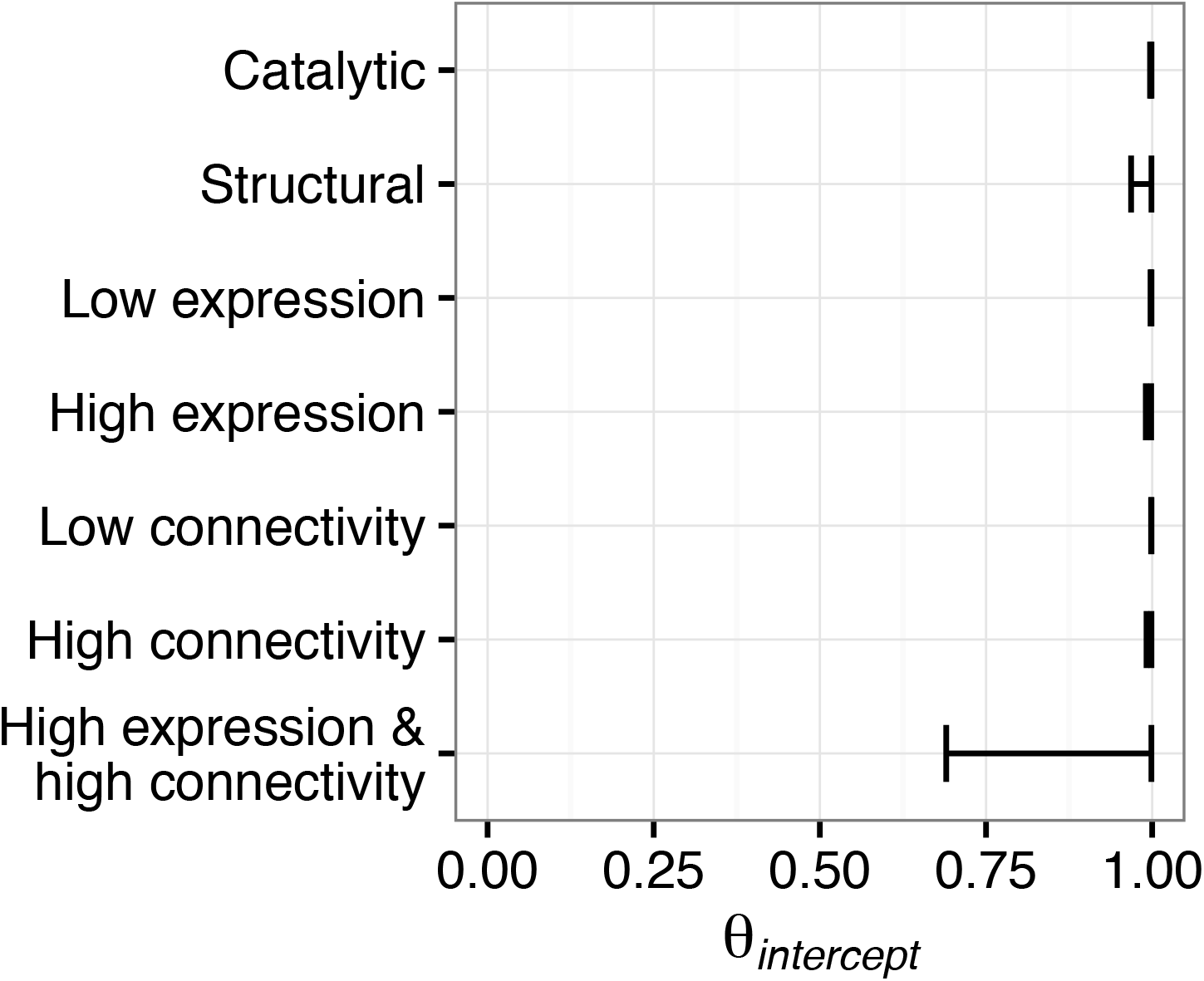
Estimation of the intercept parameter of the *h-s* relationship. Confidence interval (95%) for the estimate of *θ_intercept_* for different gene categories, combining data from *A. lyrata* and *A. thaliana* for estimation. Note that the confidence intervals for *θ_intercep_t* for different categories of genes overlap each other suggesting no difference in this parameter.

**Fig. S8.**
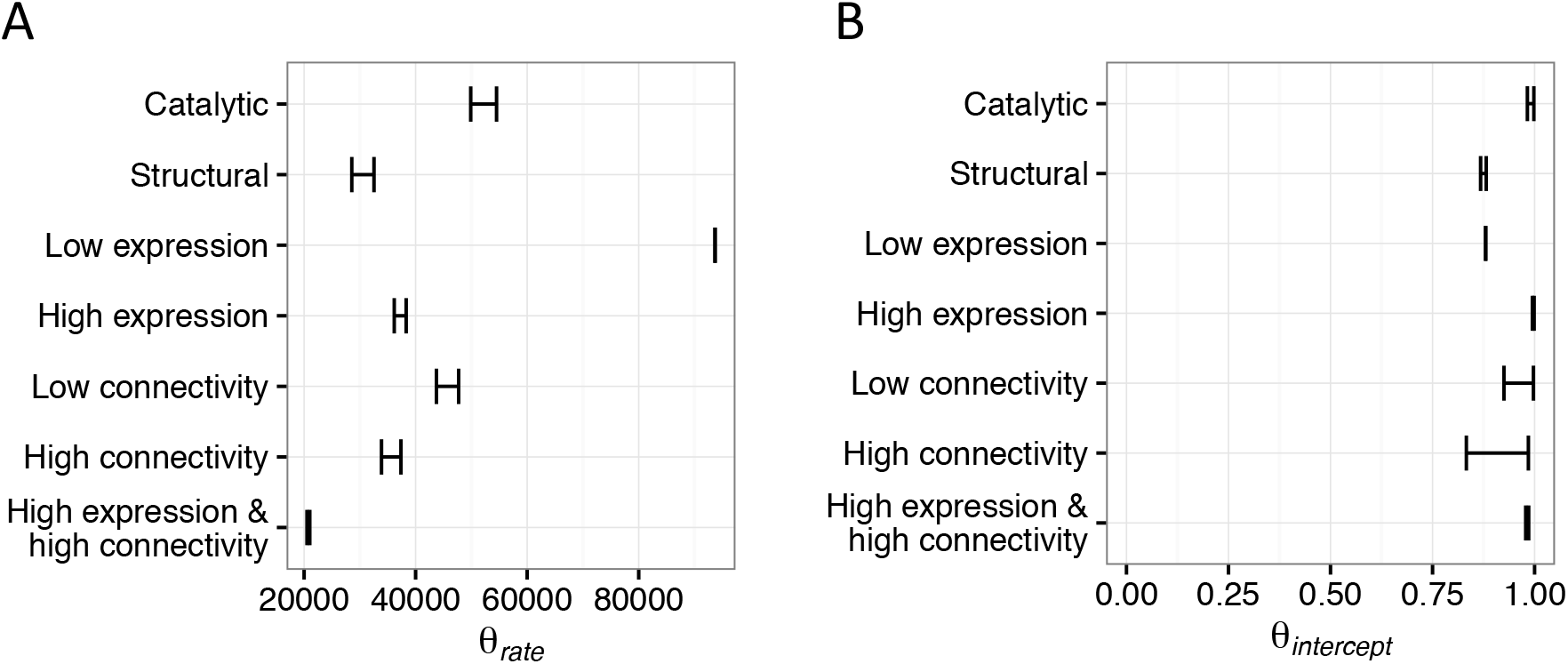
Estimation of the intercept and rate parameter of the *h-s* relationship. Confidence interval (95%) for the estimate of *θ_rate_* (A), and *θ_intercept_* (B), for different gene categories using only data from *A. lyrata* for estimation. Similar to Fig. 3C, the structural genes as well as the high expression & high connectivity genes show the smallest *θ_rate_* estimate, whereas catalytic genes, low expression genes, and low connectivity genes show the highest *θrate* estimate. However, the confidence intervals are in general larger and more variable between gene sets than when estimating the parameters using both *A. lyrata* and *A. thaliana* (Fig. 3C).

**Fig. S9.**
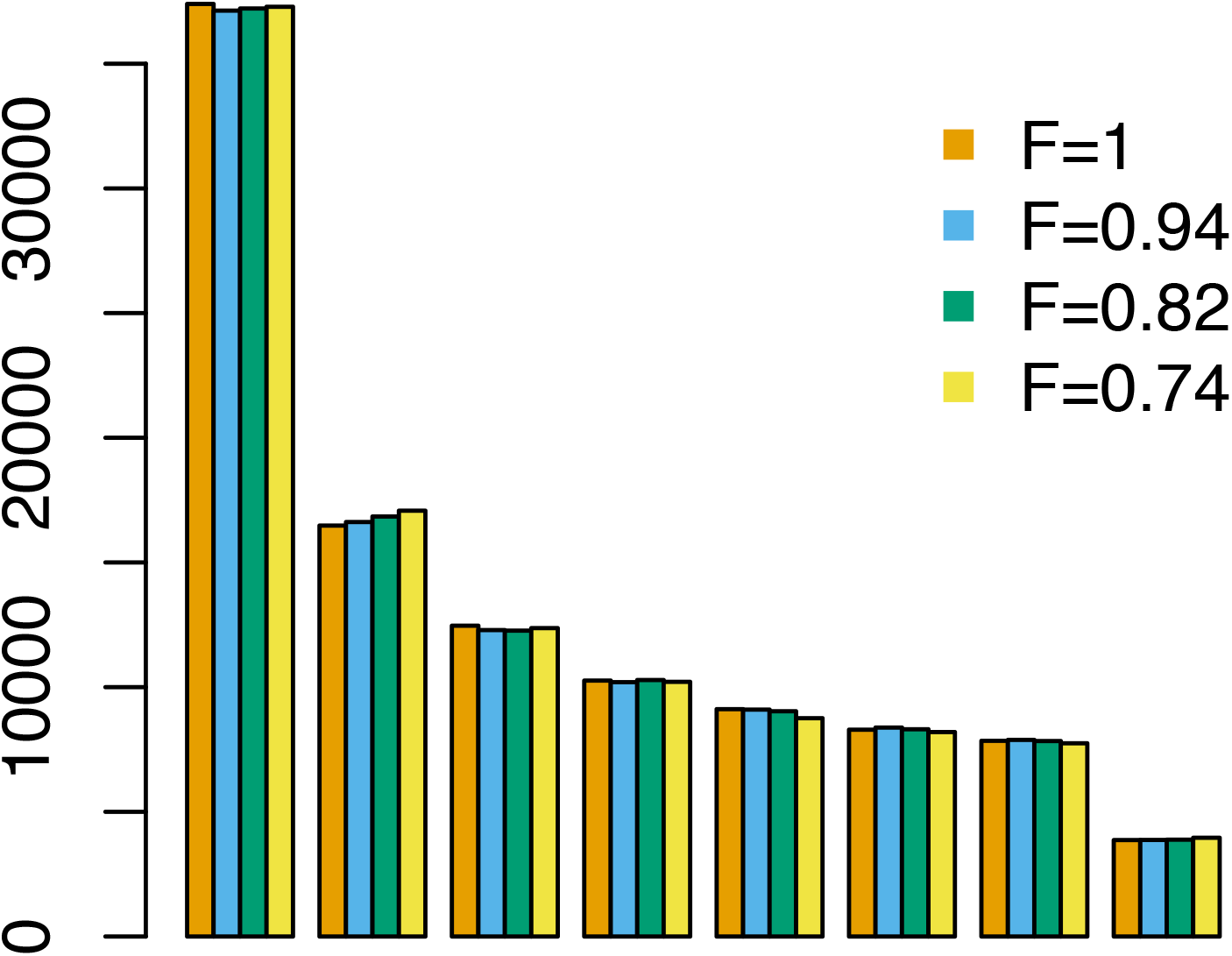
Prediction of the nonsynonymous SFS under full selfing and partial selfing. Forward simulations of the nonsynonymous SFS for *A. thaliana*, assuming either full selfing (F=1) or partial selfing with rates of 97% (F=0.94), 90% (F=0.82), or 85% (F=0.74). All four SFS agree well, suggesting that selfing in *A. thaliana* can be modeled by assuming an inbreeding coefficient of one. The simulations assume a three-epoch demographic model with parameters from Table S1. An *h-s* relationship and a gamma DFE is assumed with parameters according to the genome-wide estimates using both *A. lyrata* and *A. thaliana* (Table S3).

**Fig. S10.**
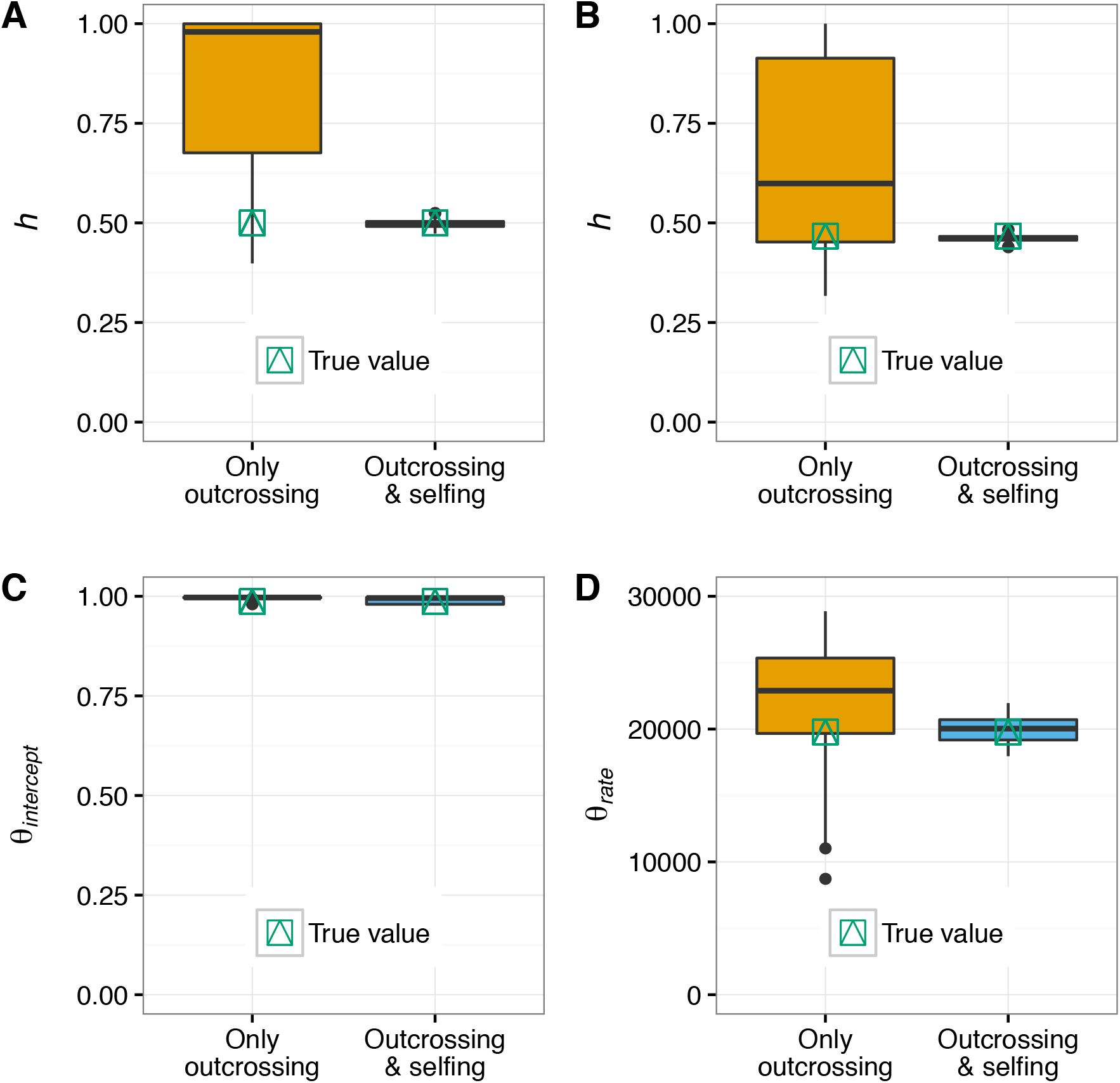
Testing the inference of dominance parameters with simulations. Data are simulated under (A) an additive model (h=0.5), (B) a constant *h* model (h=0.46), and (C, D) a *h-s* relationship model (*θ*_rate_=19773, *θ*_intercept_=0.986). True parameter values are indicated in green and MLEs from 100 replicates are shown as boxplots. Including the selfing species in the inference considerably improves estimation of the dominance parameters.

**Fig. S11.**
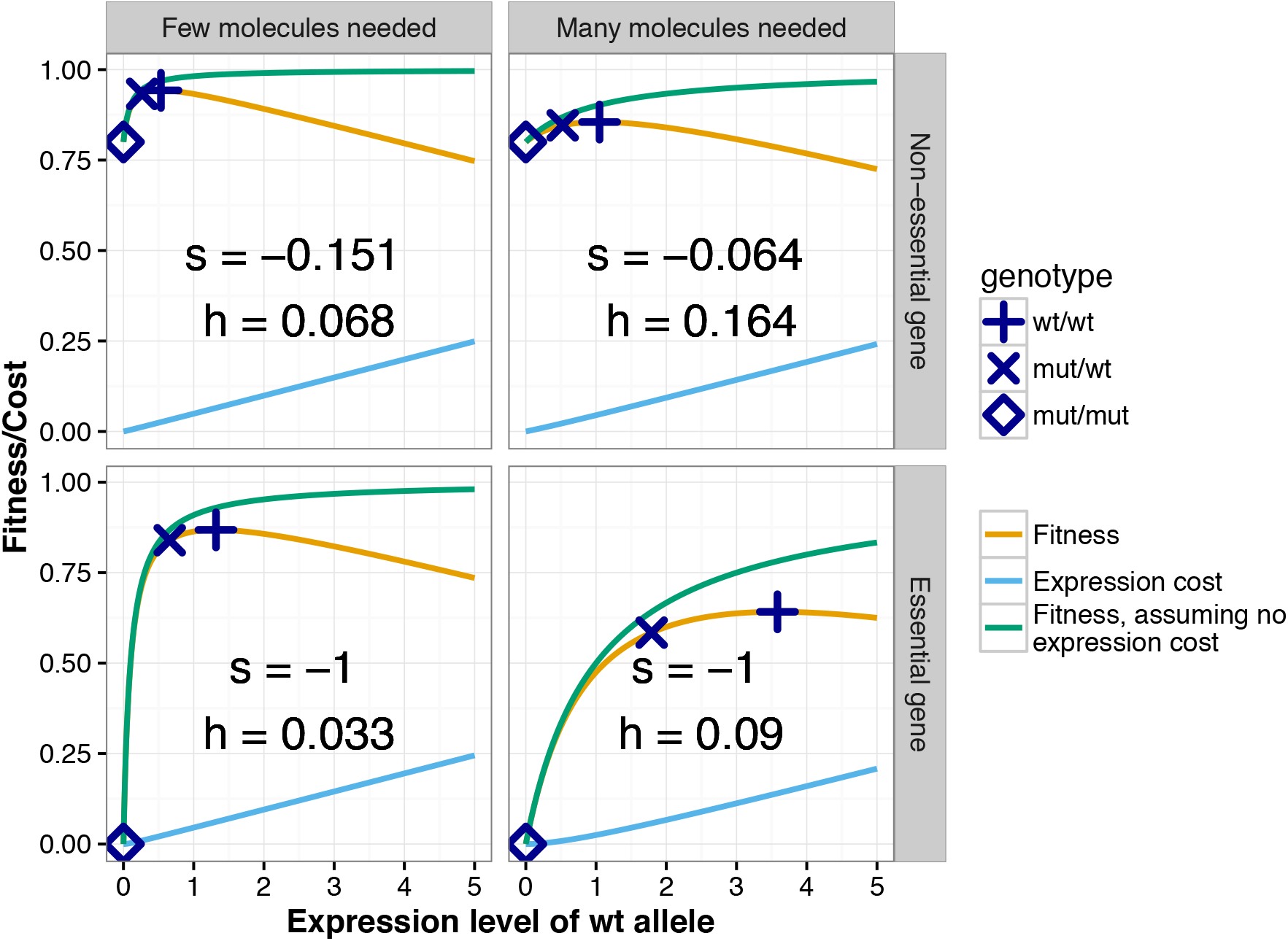
Gene expression model for the evolution of dominance. Examples of the computation of selection coefficient (s) and dominance coefficient (h) under our gene expression model for the evolution of dominance. The expression level of the homozygous wild type genotype (wt/wt) maximizes fitness after taking expression cost into account. The expression level of the gene is zero when the mutant is homozygous (mt/mt), and is half the optimal expression level when the mutant is heterozygous (wt/mt). The corresponding fitness values allow computation of s and *h* (see SI text). For non-essential genes, s and *h* are negatively related, i.e. the more deleterious mutation has a smaller *h* value. For mutations in essential genes, the gene with the higher optimal expression level has a larger *h* value than the gene with the lower optimal expression level.

**Fig. S12.**
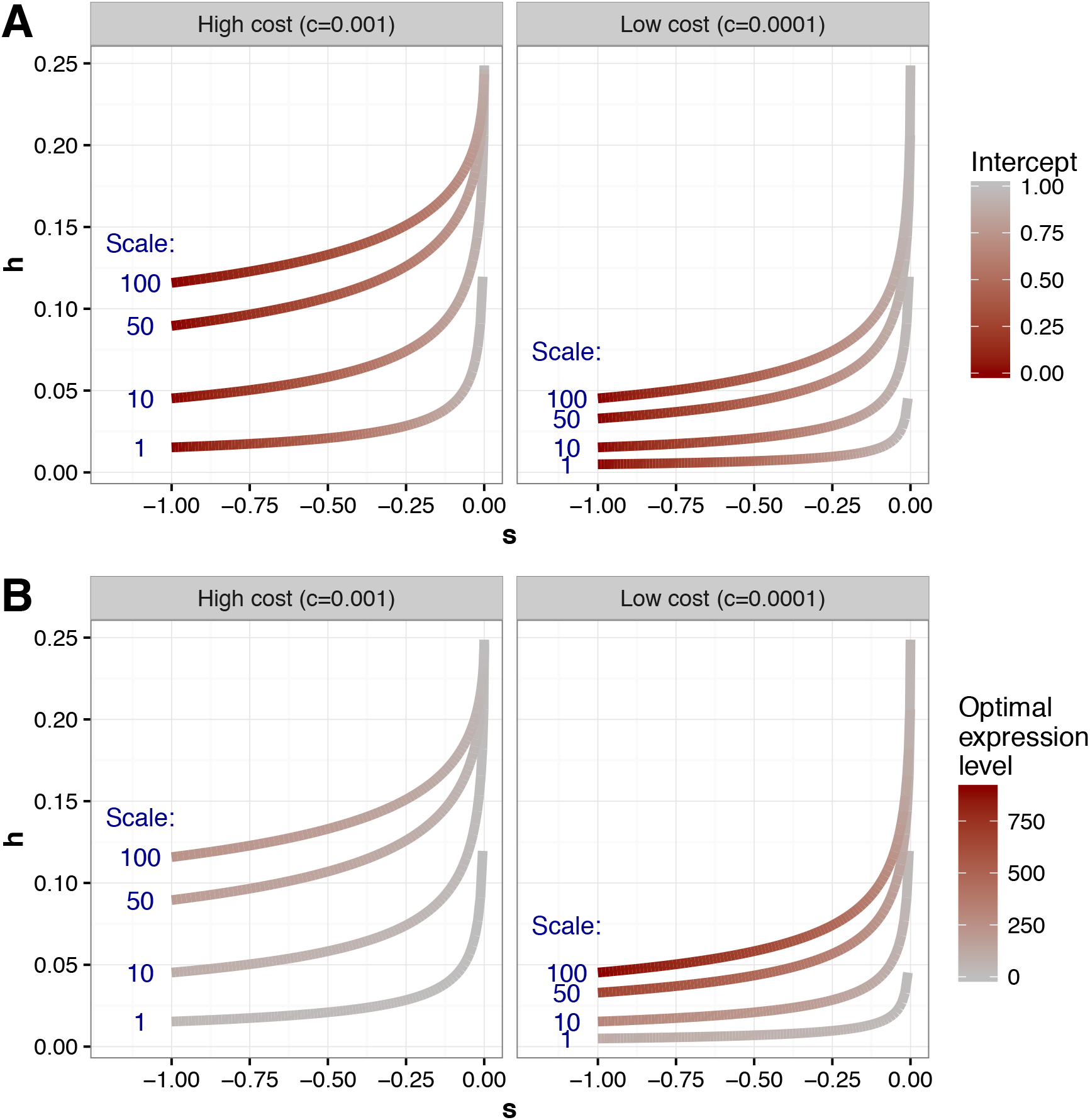
Relationship between *h* and *s* under our gene expression model for the evolution of dominance. The intercept in the model is varied continuously from 0 to 1, the scale parameter is set to 1, 10, 50, or 100, and the cost of gene expression per expression unit is set to 0.001, or 0.0001. In (A), the color scheme indicates different values of the intercept, in (B) it indicates different optimal expression levels (x_opt_).

**Table S1.** Demographic parameter estimates. Demographic parameter estimates for the two-epoch and the three-epoch model for *A. lyrata* and *A. thaliana*. The effective population size is indicated as *N_e_*, LL is the log-likelihood, and T is the time length of the epoch in generations.

| Model | Species | $N_{e,ancestral}$ | $N_{e,second\ epoch}$ | $N_{e,third\ epoch}$ | T(second epoch) | T(third epoch) | Synonymous theta | LL |
| --- | --- | --- | --- | --- | --- | --- | --- | --- |
| Two-epoch | Lyrata | 530,895 | 1,797,556 | - | 562,612 | - | 129,058 | -1095 |
|  | Thaliana | 746,148 | 100,218 | - | 568,344 | - | 199,771 | -104 |
| Three-epoch | Lyrata | 608,570 | 6,554,858 | 23,584 | 462,952 | 1,489 | 131,613 | -218 |
|  | Thaliana | 161,744 | 24,076 | 203,077 | 7,420 | 14,534 | 41,795 | -73 |

**Table S2.** Model comparison of dominance models. Likelihood ratio test statistics (Λ) and P-values when comparing different models of dominance, using only data from *A. lyrata*. The *h-s* relationship fits the data significantly better than the additive model and significantly better than a model with a single dominance coefficient.

| H0 | H1 | $\Lambda$ | P-value |
| --- | --- | --- | --- |
| Additive | Constant<br>$h \neq 0.5$ | 123 | $<1 \times 10^{-15}$ |
| Additive | $h$-$s$<br>relationship | 495 | $<1 \times 10^{-15}$ |
| Constant<br>$h \neq 0.5$ | $h$-$s$<br>relationship | 372 | $<1 \times 10^{-15}$ |

**Table S3.**
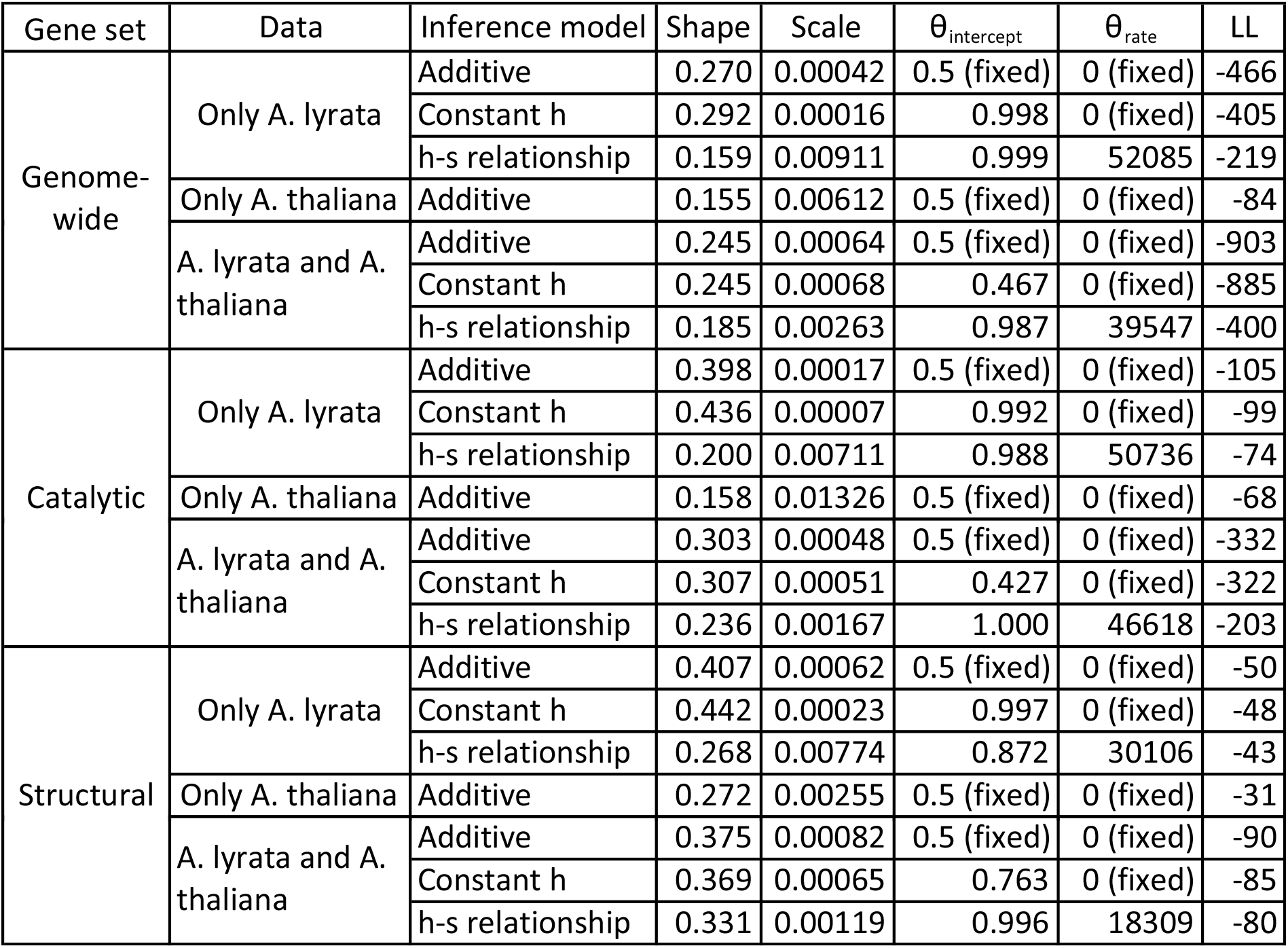
Maximum likelihood estimates of DFE and dominance parameters. Estimates for the gamma DFE parameters (shape, scale) and the two parameters of the *h-s* relationship *(*θ*_intercept_, *θ*_rate_)* for different sets of genes (Genome-wide, catalytic and structural).

**Table S4.**
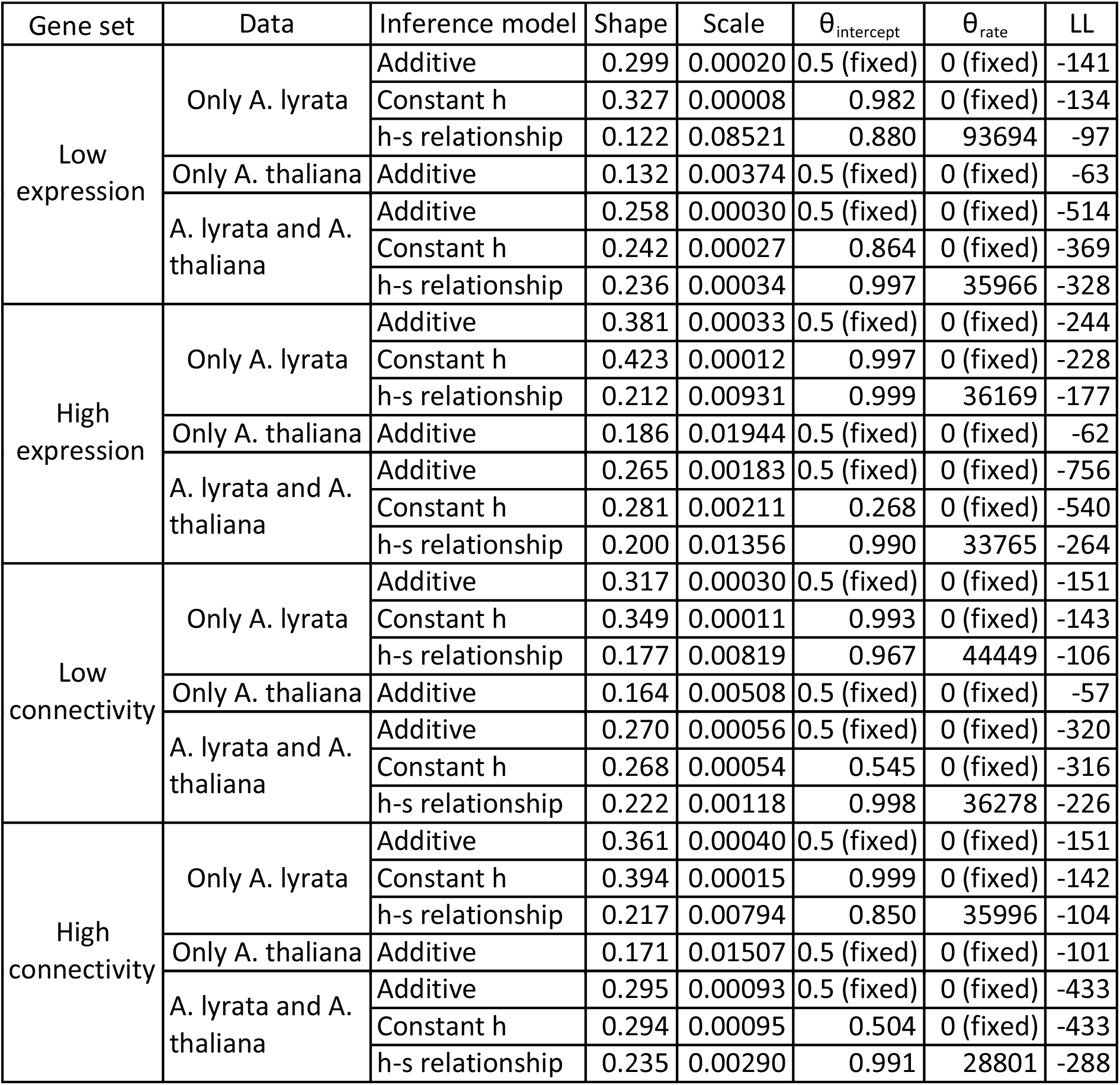
Maximum likelihood estimates of DFE and dominance parameters for different expression levels and connectivity. Estimates for the gamma DFE parameters (shape, scale) and the two parameters of the *h-s* relationship *(*θ*_intercept_, *θ*_rate_)* for different sets of genes (low and high expression level, and low and high connectivity).

| Gene set | Data | Inference model | Shape | Scale | $\theta_{\text{intercept}}$ | $\theta_{\text{rate}}$ | LL |
| --- | --- | --- | --- | --- | --- | --- | --- |
| Low expression | Only A. lyrata | Additive | 0.299 | 0.00020 | 0.5 (fixed) | 0 (fixed) | -141 |
|  |  | Constant h | 0.327 | 0.00008 | 0.982 | 0 (fixed) | -134 |
|  |  | h-s relationship | 0.122 | 0.08521 | 0.880 | 93694 | -97 |
|  | Only A. thaliana | Additive | 0.132 | 0.00374 | 0.5 (fixed) | 0 (fixed) | -63 |
|  | A. lyrata and A. thaliana | Additive | 0.258 | 0.00030 | 0.5 (fixed) | 0 (fixed) | -514 |
|  |  | Constant h | 0.242 | 0.00027 | 0.864 | 0 (fixed) | -369 |
|  |  | h-s relationship | 0.236 | 0.00034 | 0.997 | 35966 | -328 |
| High expression | Only A. lyrata | Additive | 0.381 | 0.00033 | 0.5 (fixed) | 0 (fixed) | -244 |
|  |  | Constant h | 0.423 | 0.00012 | 0.997 | 0 (fixed) | -228 |
|  |  | h-s relationship | 0.212 | 0.00931 | 0.999 | 36169 | -177 |
|  | Only A. thaliana | Additive | 0.186 | 0.01944 | 0.5 (fixed) | 0 (fixed) | -62 |
|  | A. lyrata and A. thaliana | Additive | 0.265 | 0.00183 | 0.5 (fixed) | 0 (fixed) | -756 |
|  |  | Constant h | 0.281 | 0.00211 | 0.268 | 0 (fixed) | -540 |
|  |  | h-s relationship | 0.200 | 0.01356 | 0.990 | 33765 | -264 |
| Low connectivity | Only A. lyrata | Additive | 0.317 | 0.00030 | 0.5 (fixed) | 0 (fixed) | -151 |
|  |  | Constant h | 0.349 | 0.00011 | 0.993 | 0 (fixed) | -143 |
|  |  | h-s relationship | 0.177 | 0.00819 | 0.967 | 44449 | -106 |
|  | Only A. thaliana | Additive | 0.164 | 0.00508 | 0.5 (fixed) | 0 (fixed) | -57 |
|  | A. lyrata and A. thaliana | Additive | 0.270 | 0.00056 | 0.5 (fixed) | 0 (fixed) | -320 |
|  |  | Constant h | 0.268 | 0.00054 | 0.545 | 0 (fixed) | -316 |
|  |  | h-s relationship | 0.222 | 0.00118 | 0.998 | 36278 | -226 |
| High connectivity | Only A. lyrata | Additive | 0.361 | 0.00040 | 0.5 (fixed) | 0 (fixed) | -151 |
|  |  | Constant h | 0.394 | 0.00015 | 0.999 | 0 (fixed) | -142 |
|  |  | h-s relationship | 0.217 | 0.00794 | 0.850 | 35996 | -104 |
|  | Only A. thaliana | Additive | 0.171 | 0.01507 | 0.5 (fixed) | 0 (fixed) | -101 |
|  | A. lyrata and A. thaliana | Additive | 0.295 | 0.00093 | 0.5 (fixed) | 0 (fixed) | -433 |
|  |  | Constant h | 0.294 | 0.00095 | 0.504 | 0 (fixed) | -433 |
|  |  | h-s relationship | 0.235 | 0.00290 | 0.991 | 28801 | -288 |

